# Integrin β4 promotes DNA damage-related drug resistance in triple-negative breast cancer via TNFAIP2/IQGAP1/RAC1

**DOI:** 10.1101/2023.04.17.537130

**Authors:** Huan Fang, Wenlong Ren, Qiuxia Cui, Huichun Liang, Chuanyu Yang, Wenjing Liu, Xinye Wang, Xue Liu, Yujie Shi, Jing Feng, Ceshi Chen

## Abstract

Anti-tumor drug resistance is a challenge for human triple-negative breast cancer treatment. Our previous work demonstrated that TNFAIP2 activates RAC1 to promote triple-negative breast cancer cell proliferation and migration. However, the mechanism by which TNFAIP2 activates RAC1 is unknown. In this study, we found that TNFAIP2 interacts with IQGAP1 and Integrin β4. Integrin β4 activates RAC1 through TNFAIP2 and IQGAP1 and confers DNA damage-related drug resistance in triple-negative breast cancer. These results indicate that the Integrin β4/TNFAIP2/IQGAP1/RAC1 axis provides potential therapeutic targets to overcome DNA damage-related drug resistance in triple-negative breast cancer.

## Introduction

Breast cancer is the most commonly diagnosed cancer and the leading cause of cancer death in women[1]. Although the diagnosis and treatment of breast cancer has entered the era of molecular typing, 35% of breast cancers still experience recurrence, metastasis and treatment failure[2]. According to the expression of estrogen receptor (ERα), progesterone receptor (PR) and human epidermal growth factor receptor (HER2), breast cancer is divided into ER/PR-positive, HER2-positive and triple-negative breast cancer (TNBC)[3]. For ER/PR-positive and HER2-positive breast cancer, endocrine therapies such as tamoxifen and anti-HER2 targeted therapy such as trastuzumab have achieved good efficacy. Targeted drugs for TNBC patients with BRCA1/2 mutations include two PARP inhibitors, olaparib and talazoparib. These targeted drugs cannot fully meet the clinical needs of patients with various TNBC subtypes[4]. Currently, DNA damage chemotherapy drugs, including epirubicin and cisplatin, are widely used for TNBC treatment.

TNBC often recurs and metastasizes due to the development of chemoresistance, although it is initially responsive to chemotherapeutic drugs[5]. Chemoresistance severely impacts the clinical outcomes of patients. Tumor cells become resistant to chemotherapeutic agents through several mechanisms, such as improving DNA damage repair, changing the intracellular accumulation of drugs, or increasing anti-apoptotic mechanisms[6]. Therefore, characterization of the underlying molecular mechanisms by which resistance occurs will provide opportunities to develop precise therapies to enhance the efficacy of standard chemotherapy regimens [7, 8].

TNFAIP2 is abnormally highly expressed in a variety of tumor cells, including TNBC[9], nasopharyngeal carcinoma[10], malignant glioma[11], uroepithelial carcinoma[12] and esophageal squamous cell carcinoma[13], and is associated with poor prognosis. Our previous work [9, 14]showed that TNFAIP2, as a KLF5 downstream target protein, can interact with RAC1[15], a member of the Rho small GTP enzyme family, and activate RAC1 to alter the cytoskeleton, thereby inducing filopodia and lamellipodia formation and promoting the adhesion, migration and invasion of TNBC cells. After activation, RAC1 can activate AKT, PAKs, NADPH oxidase and other related signaling pathways to promote cell survival, proliferation, adhesion, migration and invasion[16]. Activation of RAC1 can reduce the therapeutic response to trastuzumab in breast cancer and increase the resistance of TNBC cells to paclitaxel[17], but the specific mechanism of action is not completely clear.

RAC1 has been shown to play an important role in DNA damage repair. Activated RAC1 can promote the phosphorylation of the DNA damage response-related molecules ATM/ATR, CHK1/2 and H2AX by activating the activity of protein kinases such as ERK1/2, JNK and p38 [18, 19], thus improving the DNA damage repair ability and inhibiting tumor cell apoptosis[20–22]. RAC1 also promotes aldolase release and activation by changing the cytoskeleton and activates the ERK pathway to increase the pentose phosphate pathway to promote nucleic acid synthesis, providing more raw materials for DNA damage repair[23, 24]. At the same time, the interaction of RAC1 and PI3K promoted AKT phosphorylation and glucose uptake[25, 26]. Therefore, RAC1 is well established to promote the chemoresistance of breast cancer by promoting DNA damage repair.

Integrin β4 (ITGB4) is a major component of hemidesmosomes and a receptor molecule of laminin. Studies have shown that laminin-5 interacts with ITGB4 to activate RAC1 activity and promote cell migration[27] and polarization[28] by altering the cytoskeleton. Since ITGB4-positive cancer stem cell (CSC)-enriched mesenchymal cells were found to reside in an intermediate epithelial/mesenchymal phenotypic state, ITGB4 can be used to enable stratification of mesenchymal-like TNBC cells[29]. In addition, the expression ofITGB4 on ALDH^high^ breast cancer and head and neck cancer cells was significantly greater than that on ALDH^low^ cells, proving the effects that ITGB4 targets on both bulk and CSC populations[30]. Furthermore, ITGB4-overexpressing TNBC cells provided cancer-associated fibroblasts (CAFs) with ITGB4 proteins via exosomes, and ITGB4-overexpressing CAF-conditioned medium promoted the proliferation, epithelial-to-mesenchymal transition, and invasion of breast cancer cells[31]. ITGB4 also promotes breast cancer cell resistance to tamoxifen-induced apoptosis by activating the PI3K/AKT signaling pathway and promotes breast cancer cell resistance to anoikis by activating RAC1[32]. However, how ITGB4 activates RAC1 is not completely clear.

RAC1 activity is regulated by guanylate exchange factors (GEFs), GTPase activation proteins (GAPs), and guanine separation inhibitors (GDIs)[33]. GAPs typically provide the necessary catalytic groups for GTP hydrolysis, but not all GAPs function as hydrolases. IQGAP1 lacks an arginine in the GTPase binding domain and cannot exert the hydrolysis effect of GAPs[34]. IQGAP1 can increase the activity of RAC1 and CDC42[35, 36].

In this study, we demonstrated that TNFAIP2 interacts with IQGAP1 and ITGB4. ITGB4 promotes TNBC drug resistance via the TNFAIP2/IQGAP1/RAC1 axis by promoting DNA damage repair. Our results suggest that ITGB4 and TNFAIP2 might serve as promising therapeutic targets for TNBC.

## Results

### TNFAIP2 promotes TNBC DNA damage-related drug resistance

To explore the functional significance of TNFAIP2 in TNBC drug resistance, we constructed stable TNFAIP2 overexpression and TNFAIP2 knockdown HCC1806 and HCC1937 cells. As shown in Figure 1A-E, overexpression of TNFAIP2 significantly increased cell viability when treated with EPI and BMN. Additionally, knockdown of TNFAIP2 significantly decreased cell viability when treated with EPI and BMN (Figure 1F-J). We then examined the effects of TNFAIP2 on DNA damage repair and found that TNFAIP2 promotes DNA damage repair in response to EPI and BMN. TNFAIP2 overexpression decreased the protein expression levels of γH2AX, a marker of DNA damage, and cleaved-PARP, a marker of apoptosis (Figure 1K). Additionally, knockdown of TNFAIP2 significantly increased γH2AX and cleaved-PARP protein expression levels in response to EPI and BMN in both cell lines (Figure 1L).

**Figure 1.**
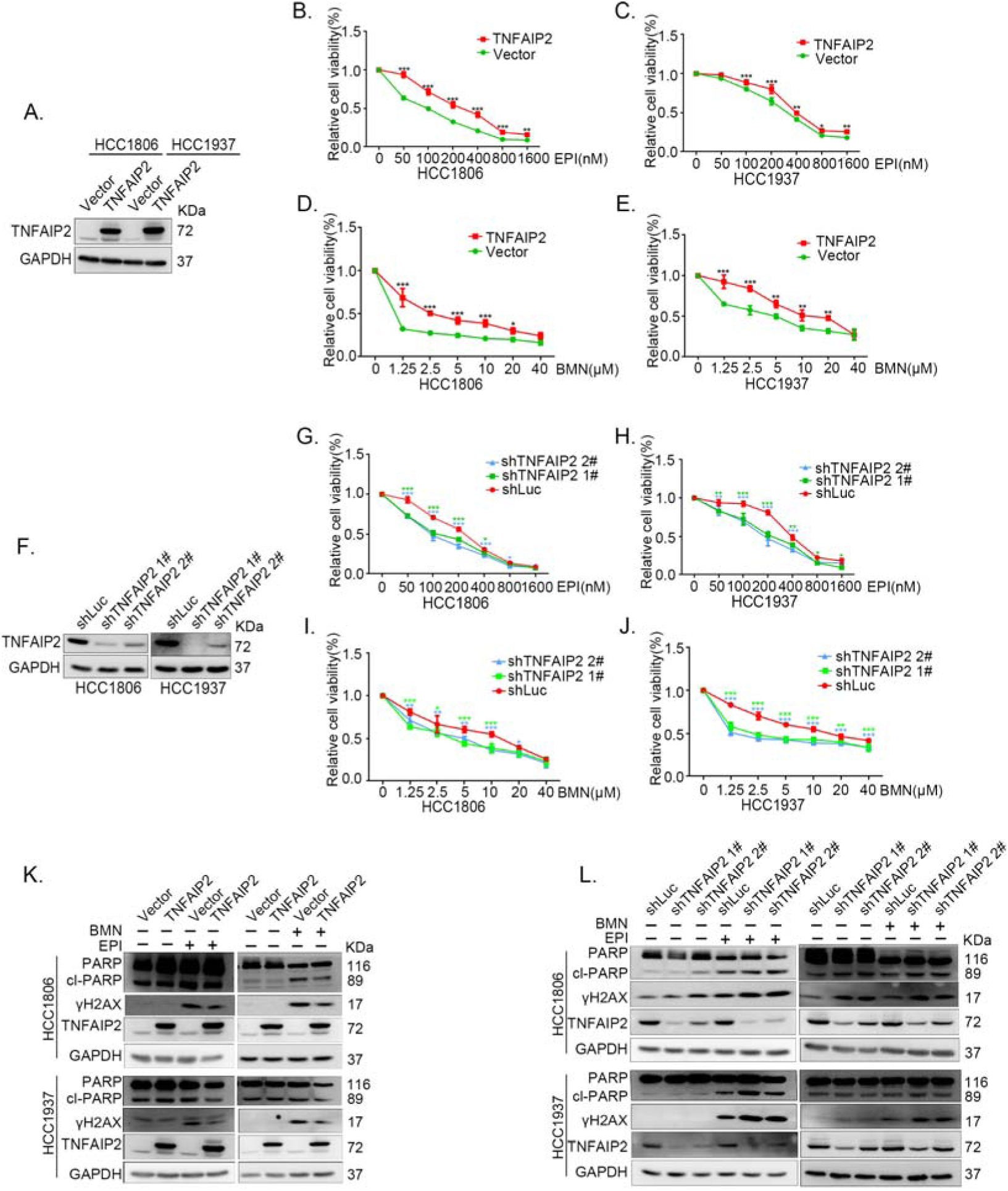
TNFAIP2 promotes TNBC DNA damage-related drug resistance. (A-E) Stable TNFAIP2 overexpression in HCC1806 and HCC1937 cells significantly increased cell viability in the presence of EPI (0-1.6 μM) or BMN (0-40 μM) treatment for 48 h, as measured by the SRB assay. Statistical analysis was performed using one-way ANOVA, n=3, * *P*<0.05, ** *P*<0.01, *** *P*<0.001. TNFAIP2 protein expression was detected by WB. (F-J) Stable TNFAIP2 knockdown in HCC1806 and HCC1937 cells significantly decreased cell viability in the presence of EPI (0-1.6 μM) or BMN (0-40 μM) treatment for 48 h, as measured by the SRB assay. Statistical analysis was performed using one-way ANOVA, n=3, * *P*<0.05, ** *P*<0.01, *** *P*<0.001. TNFAIP2 protein expression was detected by WB. (K) TNFAIP2 promoted DNA damage repair in the presence of EPI and BMN. HCC1806 and HCC1937 cells stably overexpressing TNFAIP2 were treated with 400 or 800 nM EPI for 48 h and 10 μM BMN for 24 h, respectively. TNFAIP2, γH2AX, and PARP protein expression was detected by WB. (L) TNFAIP2 knockdown increased DNA damage in the presence of EPI and BMN. Stable TNFAIP2 knockdown cells were treated with 400 or 800 nM EPI for 24 or 48h and 2.5 μM BMN for 24 h. TNFAIP2, γH2AX, and PARP protein expression was detected by WB. Figure 1-source data 1 Uncropped western blot images for Figure 1

The function of TNFAIP2 was further validated by using two other DNA damage drugs, DDP and AZD (Figure 1-figure supplement 1A-L). These results suggested that TNFAIP2 enhances TNBC cell drug resistance by promoting DNA damage repair.

### TNFAIP2 confers TNBC drug resistance *in vivo*

To test whether TNFAIP2 also decreases the sensitivity of TNBC cells to EPI and BMN *in vivo*, we performed animal experiments in nude mice. HCC1806 cells with stable TNFAIP2 knockdown were orthotopically inoculated into the fat pad of 7-week-old female mice (n=8 or 12/group). Western blotting was performed to detect the knockdown effect of TNFAIP2 protein in animal experiments (Figure 2-figure supplement 2G). When the tumor mass reached approximately 50 mm^3^, each group was divided into two subgroups to receive either EPI (2.5 mg/kg, twice a week) or vehicle control for 23 days and either BMN (1 mg/kg, twice a week) or vehicle control for 29 days. We observed that depletion of TNFAIP2 suppressed breast cancer cell growth *in vivo*. This is consistent with our previous report[9]. More importantly, TNFAIP2 depletion further decreased tumor volume when mice were treated with EPI and BMN (Figure 2A-F). Meanwhile, BMN treatment had no effect on the body weight of mice (Figure 2-figure supplement 1F). Consistently, EPI and DDP generated similar results but decreased mouse body weight due to their high toxicity (Figure 2-figure supplement 1D-E). These results suggest that inhibition of TNFAIP2 expression can overcome HCC1806 breast cancer cell drug resistance in animals.

**Figure 2.**
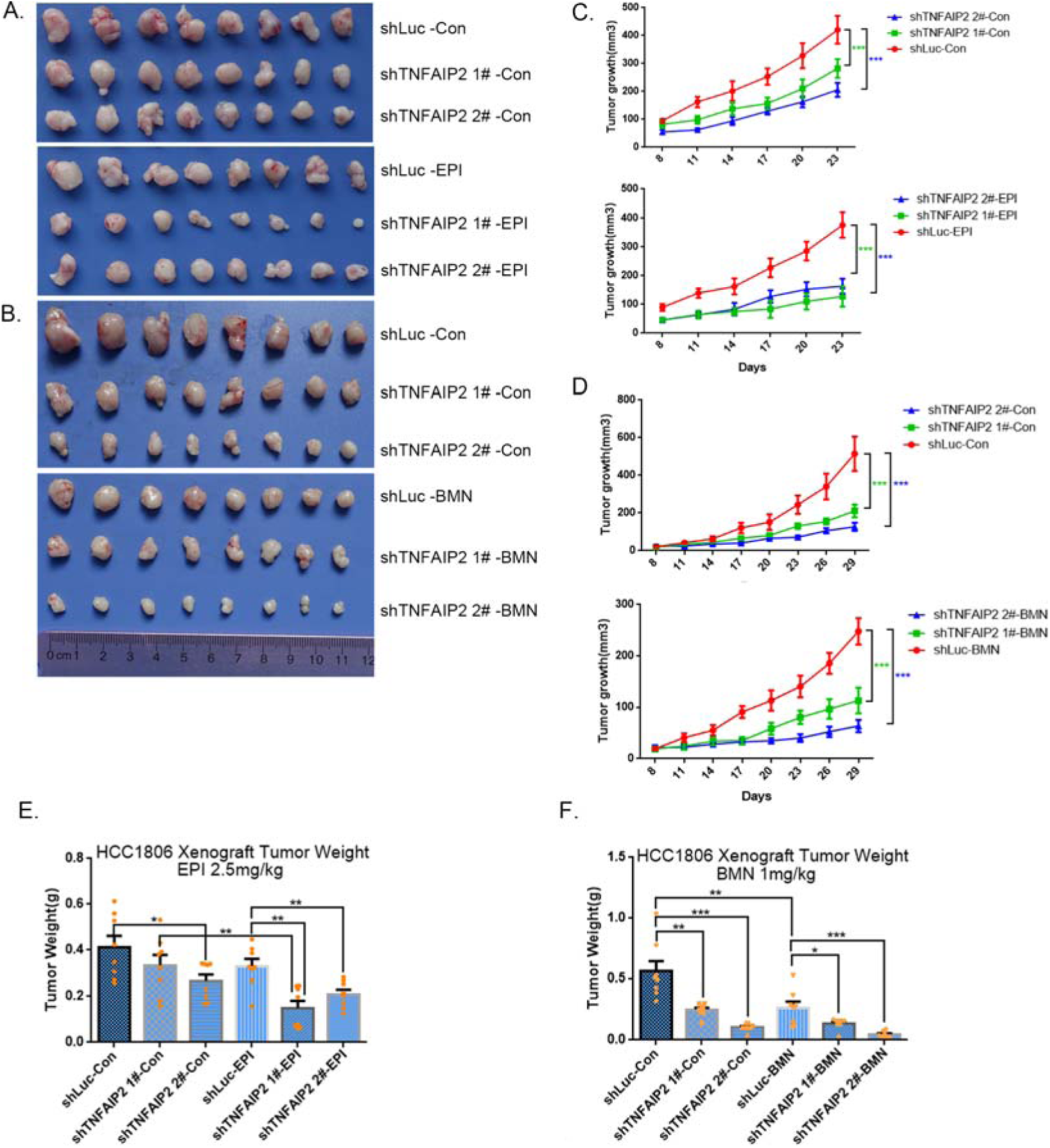
TNFAIP2 confers TNBC drug resistance *in vivo*. (A-F) TNFAIP2 knockdown increased the sensitivity ofHCC1806 breast cancer cells to EPI and BMN *in vivo*. HCC1806 cells with stable TNFAIP2 knockdown were transplanted into the fat pad of 7-week-old female nude mice. When the average tumor size reached approximately 50 mm^3^ after inoculation, mice in each group were randomly divided into two subgroups (n=4/group) to receive EPI (2.5 mg/kg), BMN (1 mg/kg) or vehicle control for 23 or 29 days (A-B). Tumor size was measured twice a week (C-D), and tumor masses were collected and weighed at the end of the experiments (E-F). *: *P*<0.05, **: *P*<0.01, ***: *P*<0.001, *t-*test.

### TNFAIP2 promotes TNBC drug resistance and DNA damage repair via RAC1

Since chemotherapeutic agents and PRAP inhibitors induce DNA damage directly or indirectly, DNA damage repair ability profoundly affects the sensitivity of cancer cells to these therapies[37, 38]. Since TNFAIP2 can activate RAC1, a well-known drug resistance protein, we investigated whether TNFAIP2 induces chemotherapeutic resistance through RAC1. We found that RAC1 knockdown abrogated the effects of TNFAIP2 overexpression-induced drug resistance to EPI and BMN in HCC1806 and HCC1937 cells (Figure 3A-F). We also found that γH2AX and cleaved-PARP protein levels were up-regulated again in RAC1 knockdown and TNFAIP2-overexpressing HCC1806 and HCC1937 cells in response to EPI and BMN (Figure 3G-J). We obtained similar results by using DDP and AZD treatment (Figure 3-figure supplement 1A-J). Collectively, these results suggest that TNFAIP2 promotes DNA damage repair and drug resistance via RAC1.

**Figure 3.**
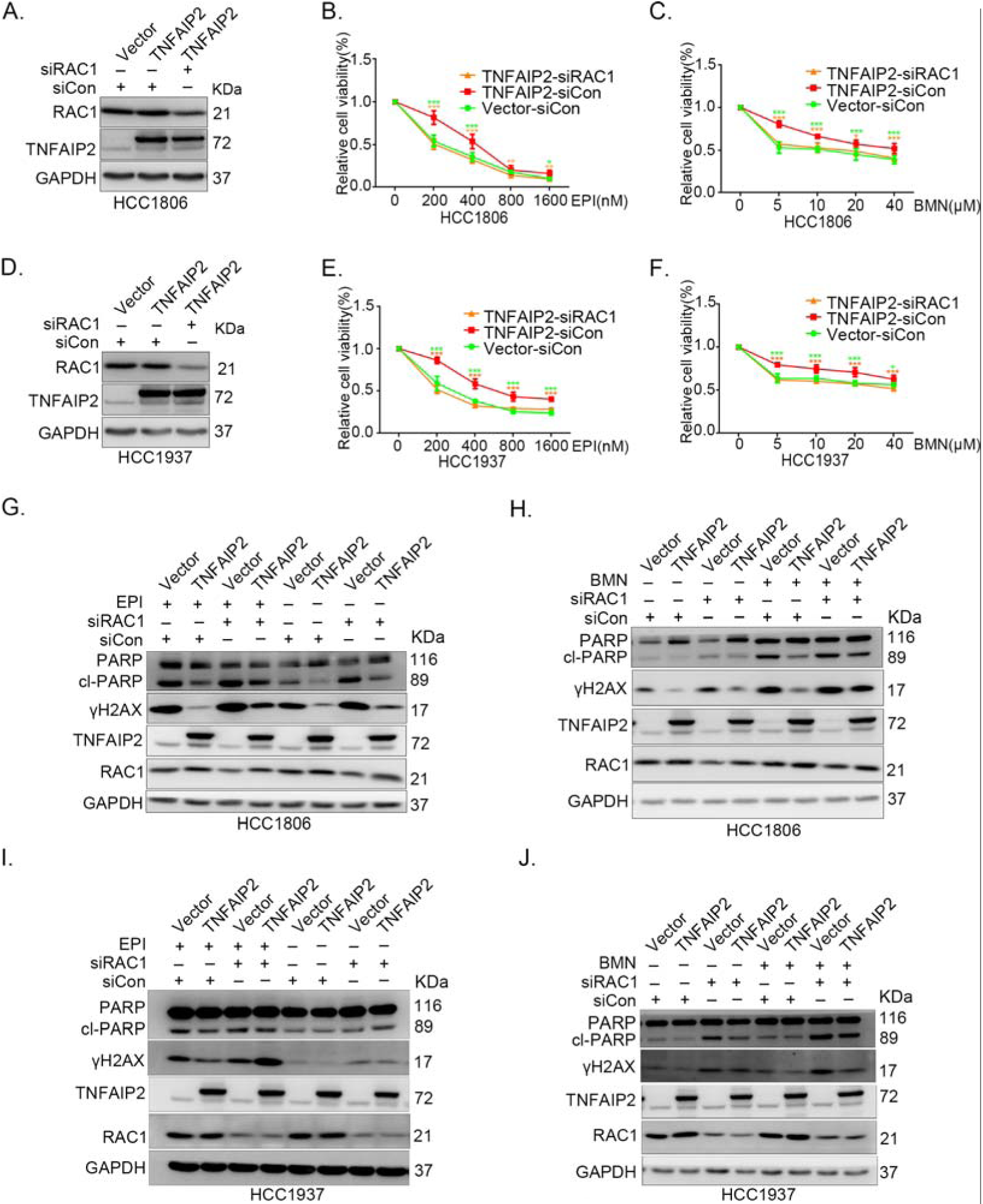
TNFAIP2 promotes TNBC drug resistance and DNA damage repair via RAC1. (A-F) RAC1 knockdown abolished TNFAIP2-induced TNBC resistance to EPI and BMN. HCC1806 (A-C) and HCC1937 (D-F) cells with stable TNFAIP2 overexpression were transfected with RAC1 or control siRNA, followed by treatment with EPI (0-1600 nM) and BMN (0-40 μM) for 48 or 72 h, respectively. Cell viability was measured by the SRB assay. Statistical analysis was performed using one-way ANOVA, n=3-9, * *P*<0.05, ** *P*<0.01, *** *P*<0.001. Protein expression levels were analyzed by WB. (G-J) RAC1 depletion abolished TNFAIP2-induced DNA damage decrease in response to EPI and BMN. HCC1806 (G-H) and HCC1937 (I-J) cells with stable TNFAIP2 overexpression were transfected with RAC1 or control siRNA, followed by treatment with EPI (400 or 800 nM) and BMN (10 μM) for 24 h, respectively. Protein expression levels were analyzed by WB. Figure 3-source data 1 Uncropped western blot images for Figure 3

### IQGAP1 mediates RAC1 activation by TNFAIP2 and promotes TNBC drug resistance

To characterize the mechanism by which TNFAIP2 activates RAC1, we performed an IP-MS experiment. We found that TNFAIP2 interacts with IQGAP1 (Figure 4A). To validate whether TNFAIP2 interacts with IQGAP1, we constructed HCC1806 cells with stable Flag-TNFAIP2 overexpression and collected Flag-tagged TNFAIP2 cell lysates for immunoprecipitation assays using Flag-M2 beads (Figure 4-figure supplement 1A).We performed immunoprecipitation using ananti-IQGAP1 antibody and found that endogenous IQGAP1 protein interacted with endogenous TNFAIP2 protein inHCC1806 cells (Figure 4B). Next, we mapped the regions of TNFAIP2 and IQGAP1 proteins responsible for the interaction by generating a series of Flag-TNFAIP2 deletion mutants and transfected them into HEK293T cells together with full-length IQGAP1. Then, we performed immunoprecipitation assays using Flag-M2 beads (Figure 4-figure supplement 1B). We demonstrated that the N-terminus (1-79 aa) of the TNFAIP2 protein interacted with IQGAP1. To explore the function of IQGAP1 in TNBC drug resistance, we knocked down IQGAP1 in HCC1806 and HCC1937 cells. As shown in Figure 4C-G, knockdown of IQGAP1 significantly decreased cell viability in the presence of EPI and BMN in both cell lines. We also examined the effects of IQGAP1 on DNA damage repair and found that IQGAP1 promotes DNA damage repair. IQGAP1 knockdown increased γH2AX and cleaved-PARP protein expression levels when HCC1806 and HCC1937 cells were treated with EPI and BMN (Figure 4H). Next, we found that IQGAP1 knockdown abrogated the effects of TNFAIP2 overexpression on resistance to EPI and BMN (Figure 4I-K, Figure 4-figure supplement 1C-E). We also found that γH2AX and cleaved-PARP protein levels were up-regulated in IQGAP1 knockdown and TNFAIP2-overexpressing HCC1806 and HCC1937 cells (Figure 4L). In addition, we found that the TNFAIP2 overexpression-induced increase in RAC1 activity was abolished by IQGAP1 knockdown (Figure 4M).

**Figure 4.**
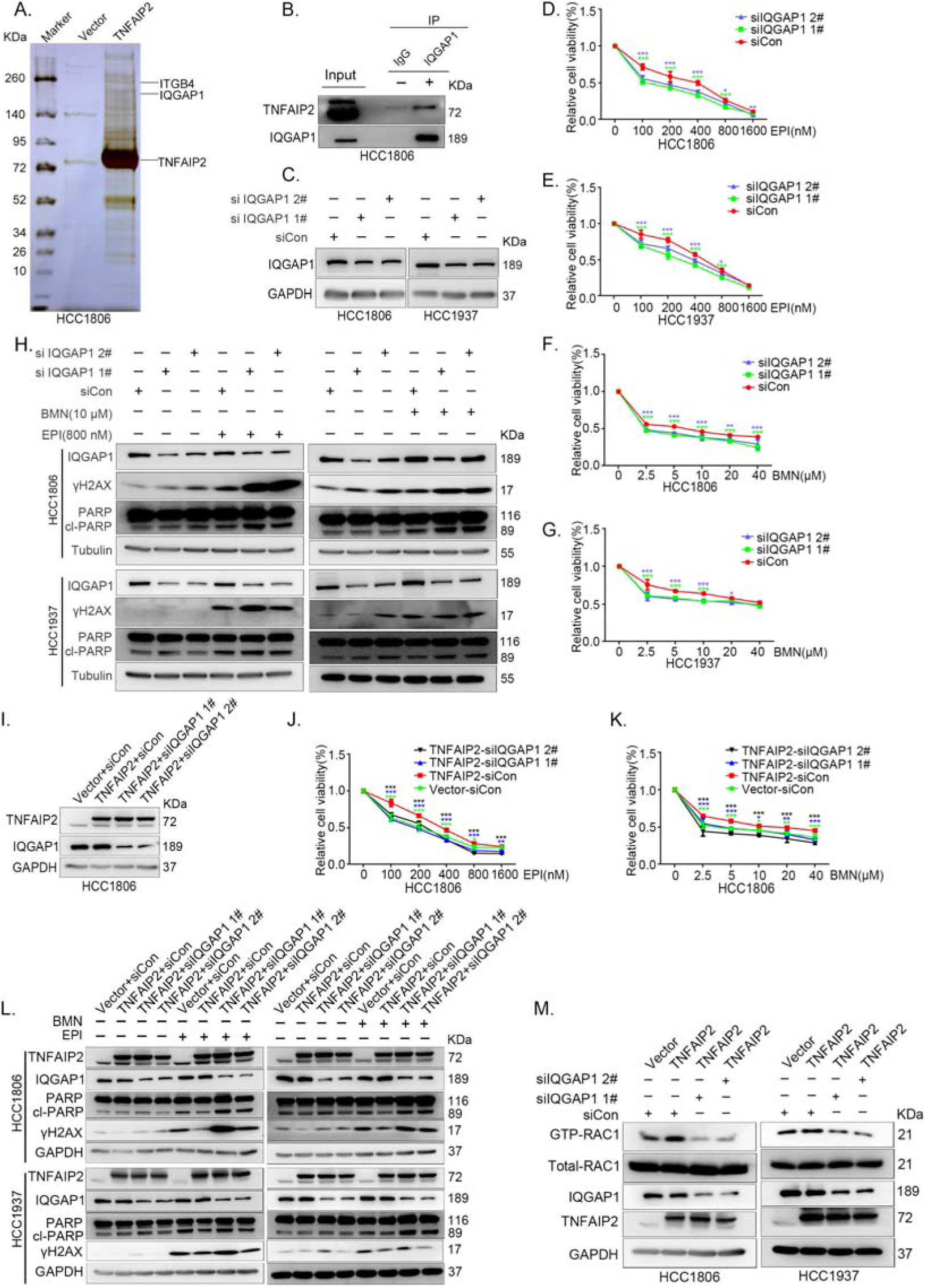
IQGAP1 mediates RAC1 activation by TNFAIP2 and promotes TNBC drug resistance. (A) The IP-MS result of TNFAIP2 in HCC1806 cells. (B) Endogenous TNFAIP2 interacts with IQGAP1 in HCC1806 cells. Endogenous TNFAIP2 protein was immunoprecipitated using ananti-IQGAP1 antibody. Immunoglobulin (Ig)G served as the negative control. Endogenous TNFAIP2 was detected by WB. (C-G) IQGAP1 knockdown in HCC1806 and HCC1937 cells significantly decreased cell viability in the presence of EPI (0-1600 nM) and BMN (0-40 μM), as measured by the SRB assay. Statistical analysis was performed using one-way ANOVA, n=3-6, * *P*<0.05, ** *P*<0.01, *** *P*<0.001. IQGAP1 protein expression was detected by WB. (H) IQGAP1 knockdown in HCC1806 and HCC1937 cells increased DNA damage of EPI and BMN. HCC1806 and HCC1937 cells with IQGAP1 knockdown were treated with 800 nM EPI for 24 h and 10 μM BMN for 24 h, respectively. ITGB4, γH2AX, and PARP protein expression was detected by WB. (I-K) IQGAP1 knockdown abolished TNFAIP2-confered resistance to EPI and BMN. HCC1806 cells with stable TNFAIP2 overexpression were transfected with IQGAP1 or control siRNA, followed by treatment with EPI (0-1600 nM) and BMN (0-40 μM) for 48 or 72 h, respectively. Cell viability was measured by the SRB assay. Statistical analysis was performed using one-way ANOVA, n=3, * *P*<0.05, ** *P*<0.01, *** *P*<0.001. IQGAP1 protein expression was detected by WB. (L) IQGAP1 knockdown abolished TNFAIP2-confered resistance to EPI and BMN. HCC1806 and HCC1937 cells with stable TNFAIP2 overexpression were transfected with IQGAP1 or control siRNA, followed by treatment with EPI (800 nM) and BMN (10 μM) for 24 h, respectively. Protein expression levels were analyzed by WB. (M) IQGAP1 knockdown abolished TNFAIP2-confered RAC1 activation. HCC1806 and HCC1937 cells with stable TNFAIP2 overexpression were transfected with IQGAP1 or control siRNA. GTP-RAC1 levels were assessed using PAK-PBD beads. Figure 4-source data 1 Uncropped western blot images for Figure 4

### ITGB4 interacts with TNFAIP2 and promotes TNBC drug resistance and DNA damage repair

In addition to IQGAP1, TNFAIP2 may interact with ITGB4 (Figure 4A). To validate whether TNFAIP2 interacts with ITGB4, we immunoprecipitated exogenous Flag-tagged TNFAIP2 proteins from HCC1806 cells by using Flag-M2 beads and detected endogenous ITGB4 proteins (Figure 5A).To further confirm the protein protein interaction between endogenous TNFAIP2 and ITGB4 proteins, we collected HCC1806 cell lysates and performed immunoprecipitation using an anti-TNFAIP2/ITGB4 antibody and found that endogenous TNFAIP2/ITGB4 protein interacted with endogenous ITGB4/TNFAIP2 protein (Figure 5B-C). We further mapped the regions of TNFAIP2 and ITGB4 proteins responsible for the interaction (Figure 5-figure supplement 2I) by generating a series of Flag-TNFAIP2/GST-fused TNFAIP2 deletion mutants and transfected them into HEK293T cells together with full-length GST-fused ITGB4/ITGB4. Then, we performed immunoprecipitation assays using Flag-M2 beads and glutathione beads. As shown in Figure 5-figure supplement 2J-K, the N-terminus (218-287aa) of the TNFAIP2 (TNFAIP2-S-N1-3) protein interacted with ITGB4. To map the domains of ITGB4 that interact with TNFAIP2, we transfected Flag-tagged full-length TNFAIP2 into HEK293T cells with full-length or truncated ITGB4. We found that the C-terminus (710-740 aa) of the ITGB4 protein interacted with TNFAIP2 (Figure 5-figure supplement 2L-M). Taken together, these results suggest that TNFAIP2 interacts with ITGB4 and that their interaction is mediated through the N-terminus of TNFAIP2 and the C-terminus of ITGB4.

**Figure 5.**
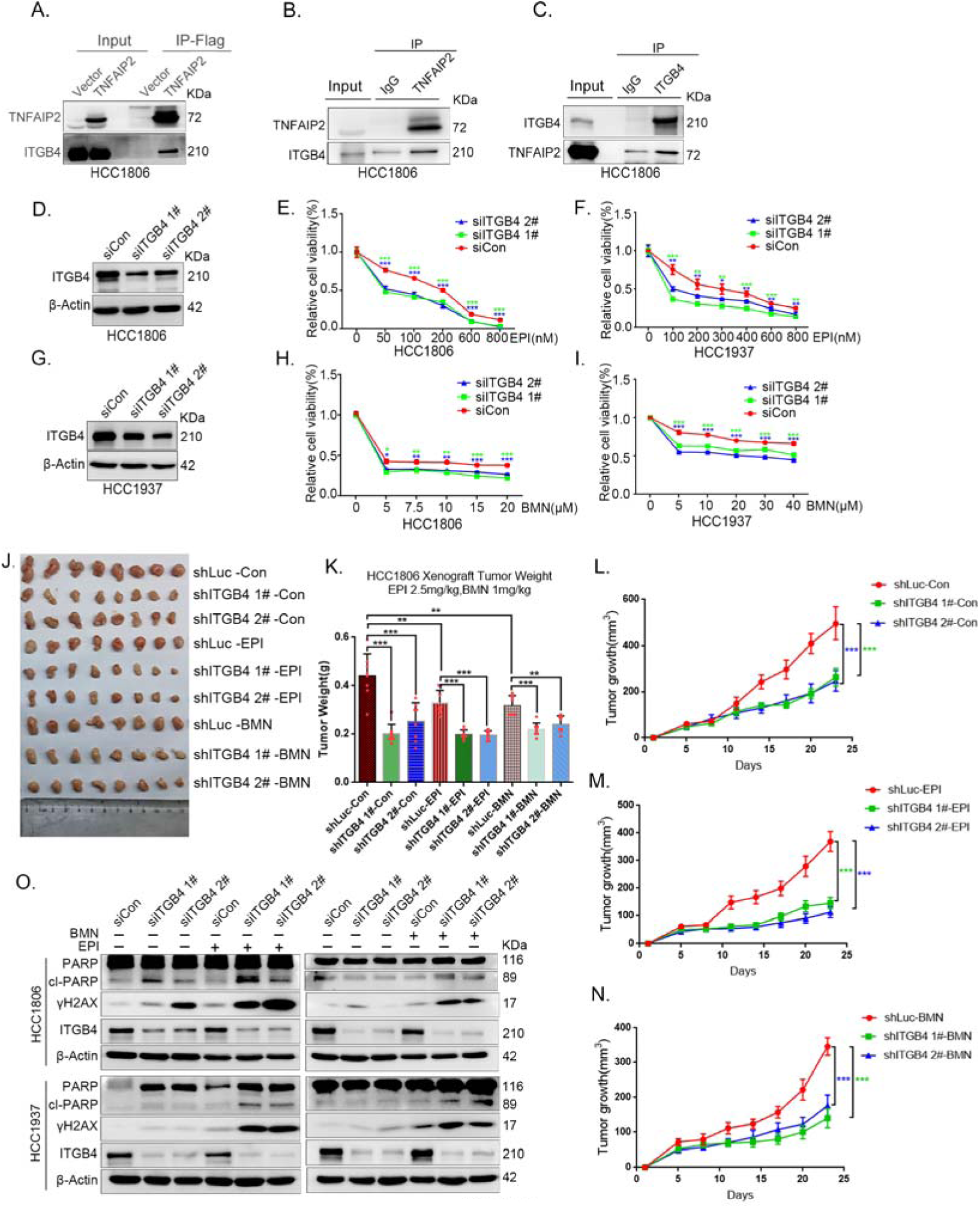
ITGB4 interacts with TNFAIP2 and promotes TNBC drug resistance and DNA damage repair. (A) TNFAIP2 interacts with ITGB4.HCC1806 cells with stable TNFAIP2 overexpression were collected from flag-tagged TNFAIP2 cell lysates for immunoprecipitation assays using Flag-M2 beads, and ITGB4 was detected by WB. (B) Endogenous TNFAIP2 interacts with ITGB4 in HCC1806 cells. Endogenous TNFAIP2 protein was immunoprecipitated using an anti-TNFAIP2 antibody. IgG served as the negative control. Endogenous ITGB4 was detected by WB. (C) Endogenous ITGB4 interacts with TNFAIP2 in HCC1806 cells. Endogenous ITGB4 protein was immunoprecipitated using an anti-ITGB4 antibody. IgG served as the negative control. Endogenous TNFAIP2 was detected by WB. (D-I) ITGB4 knockdown in HCC1806 and HCC1937 cells significantly decreased cell viability in the presence of EPI (0-800 nM) and BMN (0-40 μM), as measured by the SRB assay. Statistical analysis was performed using one-way ANOVA, n=3, * *P*<0.05, ** *P*<0.01, *** *P*<0.001. ITGB4 protein expression was detected by WB. (J-N) ITGB4 depletion promotes HCC1806 breast cancer cell sensitivity to EPI and BMN treatment *in vivo*. HCC1806 cells with stable ITGB4 knockdown were transplanted into the fat pad of 7-week-old female nude mice. When the average tumor size reached approximately 50 mm^3^ after inoculation, the mice in each group were randomly divided into two subgroups (n=4/group) to receive EPI (2.5 mg/kg), BMN (1 mg/kg) or vehicle control for 22 days (J). Tumor masses were collected and weighed at the end of the experiments (K), and tumor size was measured twice a week (L-N).*: *P*<0.05, **: *P*<0.01, ***: *P*<0.001, *t*-test. (O) ITGB4 knockdown increased DNA damage of EPI and BMN. HCC1806 and HCC1937 cells with ITGB4 knockdown were treated with 400 nM EPI for 24 h and 5 μM BMN for 24 h, respectively. ITGB4, γH2AX, and PARP protein expression was detected by WB. Figure 5-source data 1 Uncropped western blot images for Figure 5

To explore the function of ITGB4 in TNBC drug resistance, we knocked down ITGB4 in HCC1806 and HCC1937 cells. As shown in Figure 5D-I, knockdown of ITGB4 significantly decreased cell viability in the presence of EPI and BMN in both cell lines. Knockdown of ITGB4 also suppressed HCC1806 xenograft growth *in vivo*. The knockdown effect of ITGB4 protein in animal experiments was confirmed by Western blotting (Figure 5-figure supplement 2N). More importantly, ITGB4 knockdown further decreased tumor volume when mice were treated with EPI and BMN (Figure 5J-N). Meanwhile, BMN treatment had no effect on the body weight of mice, but EPI treatment decreased mouse body weight due to its toxicity (Figure 5-figure supplement 1H). We then examined the effects of ITGB4 on DNA damage repair and found that ITGB4 promotes DNA damage repair in response to EPI and BMN. ITGB4 knockdown increased γH2AX and cleaved-PARP protein expression levels when HCC1806 and HCC1937 cells were treated with EPI and BMN (Figure 5O). Furthermore, the function of ITGB4 was validated by using two other drugs, DDP and AZD (Figure 5-figure supplement 1A-G). These results suggested that ITGB4 increases TNBC drug resistance and promotes DNA damage repair.

### ITGB4 activates RAC1 through TNFAIP2 and IQGAP1

It is well known that ITGB4 can activate RAC1[27] and that TNFAIP2 interacts with RAC1 and activates it[9]. To test whether ITGB4 activates RAC1 through TNFAIP2, we measured the levels of GTP-bound RAC1 in ITGB4-overexpressing and ITGB4-knockdown cells. Overexpression of ITGB4 significantly increased the levels of GTP-bound RAC1 in both HCC1806 and HCC1937 cells (Figure 6A). In agreement with this observation, knockdown of ITGB4 significantly decreased the levels of GTP-bound RAC1 in both cell lines (Figure 6B). Next, we knocked down TNFAIP2 in ITGB4-overexpressing HCC1806 and HCC1937 cells and found that ITGB4-increased RAC1 activity was blocked by TNFAIP2 knockdown (Figure 6C-D). Collectively, these results demonstrate that ITGB4 activates RAC1 through TNFAIP2.

**Figure 6.**
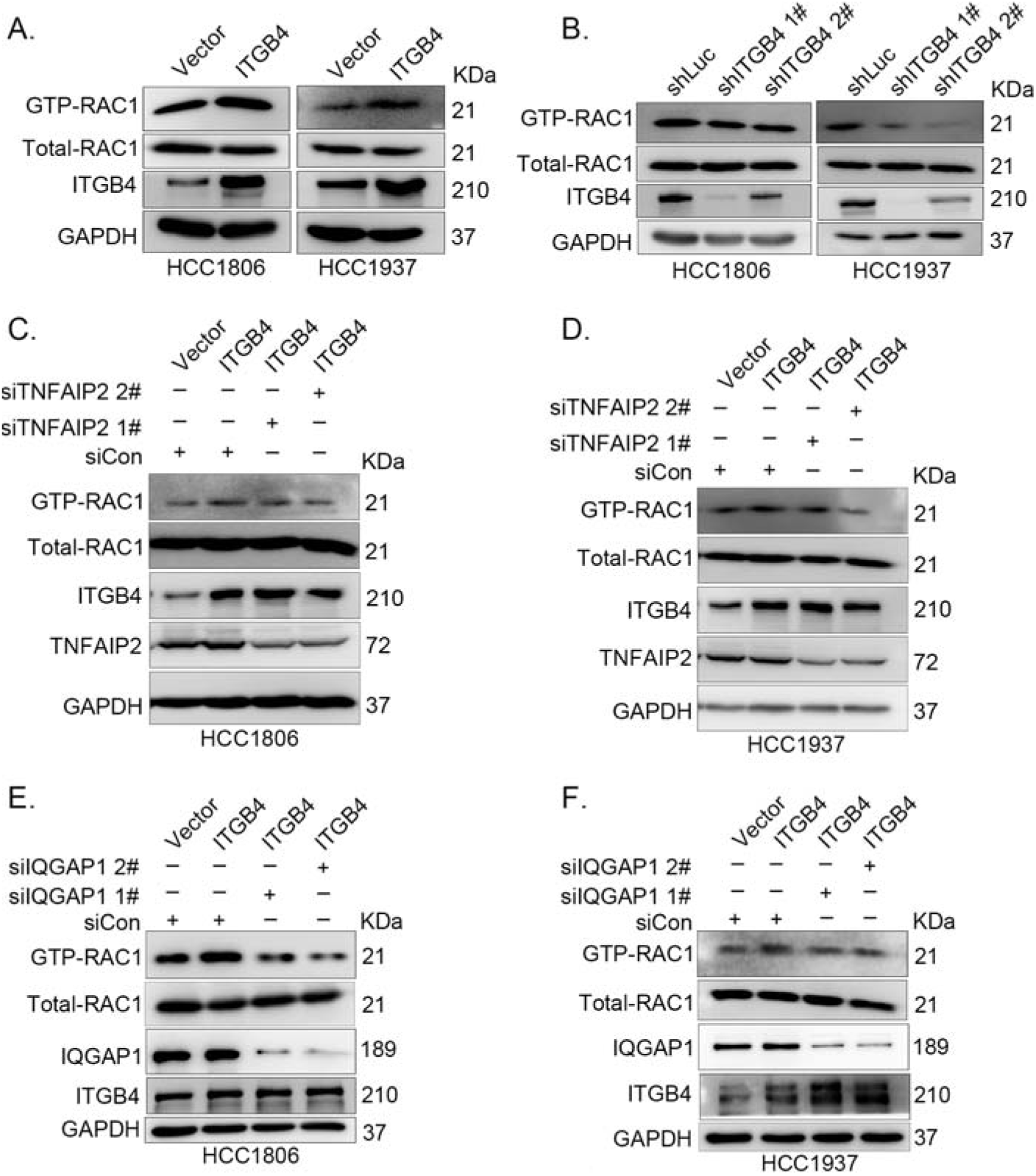
ITGB4 activates RAC1 through TNFAIP2 and IQGAP1. (A) Overexpression of ITGB4 increased GTP-RAC1 levels in HCC1806 and HCC1937 cells. GTP-RAC1 were assessed using PAK-PBD beads. (B) Knockdown of ITGB4 by shRNA decreased GTP-RAC1 levels in HCC1806 and HCC1937 cells. (C-D) ITGB4 activates RAC1 through TNFAIP2. HCC1806 (C) and HCC1937 (D) cells with stable ITGB4 overexpression were transfected with TNFAIP2 or control siRNA. (E-F) ITGB4 activates RAC1 through IQGAP1. HCC1806 (E) and HCC1937 (F) cells with stable ITGB4 overexpression were transfected with IQGAP1 or control siRNA. Figure 6-source data 1 Uncropped western blot images for Figure 6

It has been reported that RAC1 activity is promoted by IQGAP1[34] and that TNFAIP2 activates RAC1 through IQGAP1 (Figure 4P). We wondered whether ITGB4 activates RAC1 through IQGAP1; therefore, we knocked down IQGAP1 in HCC1806 and HCC1937 cells with stable overexpression of ITGB4 and found that the ITGB4-induced increase in RAC1 activity was abolished by IQGAP1 knockdown (Figure 6E-F). These results suggest that ITGB4 activates RAC1 through TNFAIP2 and IQGAP1.

### ITGB4 promotes TNBC drug resistance via TNFAIP2/IQGAP1/RAC1

Since ITGB4, TNFAIP2, and IQGAP1 promote drug resistance by promoting DNA damage repair in TNBC, we wondered whether ITGB4 promoted drug resistance through the TNFAIP2/IQGAP1/RAC1 axis. We knocked down TNFAIP2, IQGAP1, and RAC1 in ITGB4-overexpressing cells and found that blocking the TNFAIP2/IQGAP1/RAC1 axis increased the sensitivity of ITGB4-overexpressing HCC1806 (Figure 7A-I) and HCC1937 cells to EPI and BMN (Figure 7-figure supplement 2O-W). We also found that γH2AX and cleaved-PARP levels were upregulated in TNFAIP2/IQGAP1/RAC1 knockdown HCC1806 and HCC1937 cells stably expressing ITGB4 in the presence of EPI and BMN (Figure 7J-L, Figure 7-figure supplement 2X-Z). DDP and AZD treatment generated similar results (Figure 7-figure supplement 1A-N). Together, these results suggest that ITGB4 promotes DNA damage repair and drug resistance via the TNFAIP2/IQGAP1/RAC1 axis.

**Figure 7.**
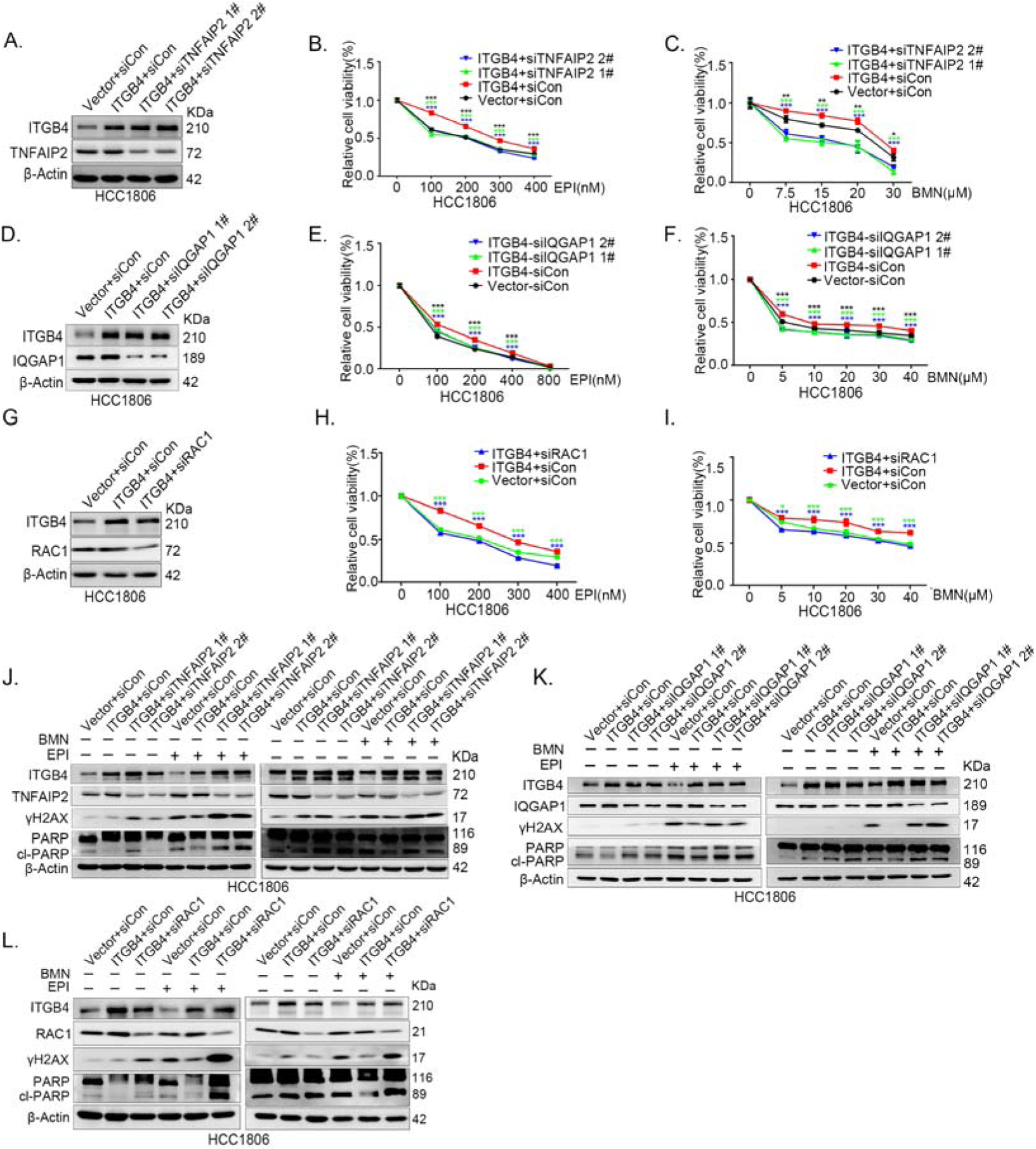
ITGB4 promotes TNBC drug resistance via TNFAIP2/IQGAP1/RAC1. (A-C) ITGB4 promotes TNBC drug resistance through TNFAIP2. TNFAIP2 knockdown abolished ITGB4-induced resistance to EPI and BMN. HCC1806 cells with stable ITGB4 overexpression were transfected with TNFAIP2 or control siRNA, followed by treatment with EPI (0-400 nM) and BMN (0-30 μM) for 48 or 72 h, respectively. Cell viability was measured by the SRB assay. Statistical analysis was performed using one-way ANOVA, n=3, * *P*<0.05, ** *P*<0.01, *** *P*<0.001. Protein expression levels were analyzed by WB. (D-F) ITGB4 promotes TNBC drug resistance through IQGAP1. HCC1806 cells with stable ITGB4 overexpression were transfected with IQGAP1 or control siRNA, followed by treatment with EPI (0-800 nM) and BMN (0-40 μM) for 48 or 72 h, respectively. Cell viability was measured by the SRB assay. Statistical analysis was performed using one-way ANOVA, n=3, * *P*<0.05, ** *P*<0.01, *** *P*<0.001. Protein expression levels were analyzed by WB. (G-I) ITGB4 promotes TNBC drug resistance through RAC1. HCC1806 cells with stable ITGB4 overexpression were transfected with RAC1 or control siRNA, followed by treatment with EPI (0-400 nM) and BMN (0-40 μM) for 48 or 72 h, respectively. Cell viability was measured by the SRB assay. Statistical analysis was performed using one-way ANOVA, n=3, * *P*<0.05, ** *P*<0.01, *** *P*<0.001. Protein expression levels were analyzed by WB. (J) ITGB4 promotes DNA damage repair through TNFAIP2. HCC1806 cells with stable ITGB4 overexpression were transfected with TNFAIP2 or control siRNA, followed by treatment with EPI (400 nM) and BMN (5 μM) for 24 h. Protein expression levels were analyzed by WB. (K) ITGB4 promotes DNA damage repair through IQGAP1. HCC1806 cells with stable ITGB4 overexpression were transfected with IQGAP1 or control siRNA, followed by treatment with EPI (400 nM) and BMN (5 μM) for 24 h. Protein expression levels were analyzed by WB. (L) ITGB4 promotes DNA damage repair through RAC1. HCC1806 cells with stable ITGB4 overexpression were transfected with RAC1 or control siRNA, followed by treatment with EPI (400 nM) and BMN (5 μM) for 24 h. Protein expression levels were analyzed by WB. Figure 7-source data 1 Uncropped western blot images for Figure 7

### TNFAIP2 expression levels positively correlated with ITGB4 in TNBC tissues

To test whether ITGB4 and TNFAIP2 are co-expressed in TNBC, we collected 135 TNBC specimens for IHC (the IQGAP1 antibody did not work for IHC). Clinical pathological parameters, including patient age, tumor size, lymph node status at the time of diagnosis, and follow-up status, including adjuvant treatment and tumor recurrence, were retrospectively obtained from the Department of Pathology, Henan Provincial People’s Hospital, Zhengzhou University, China. We performed IHC analyses on two breast cancer tissue chips containing a total of 135 patients with TNBC (Figure 8A-D). TNFAIP2 and ITGB4 protein expression levels were significantly positively correlated (Figure 8E).

**Figure 8.**
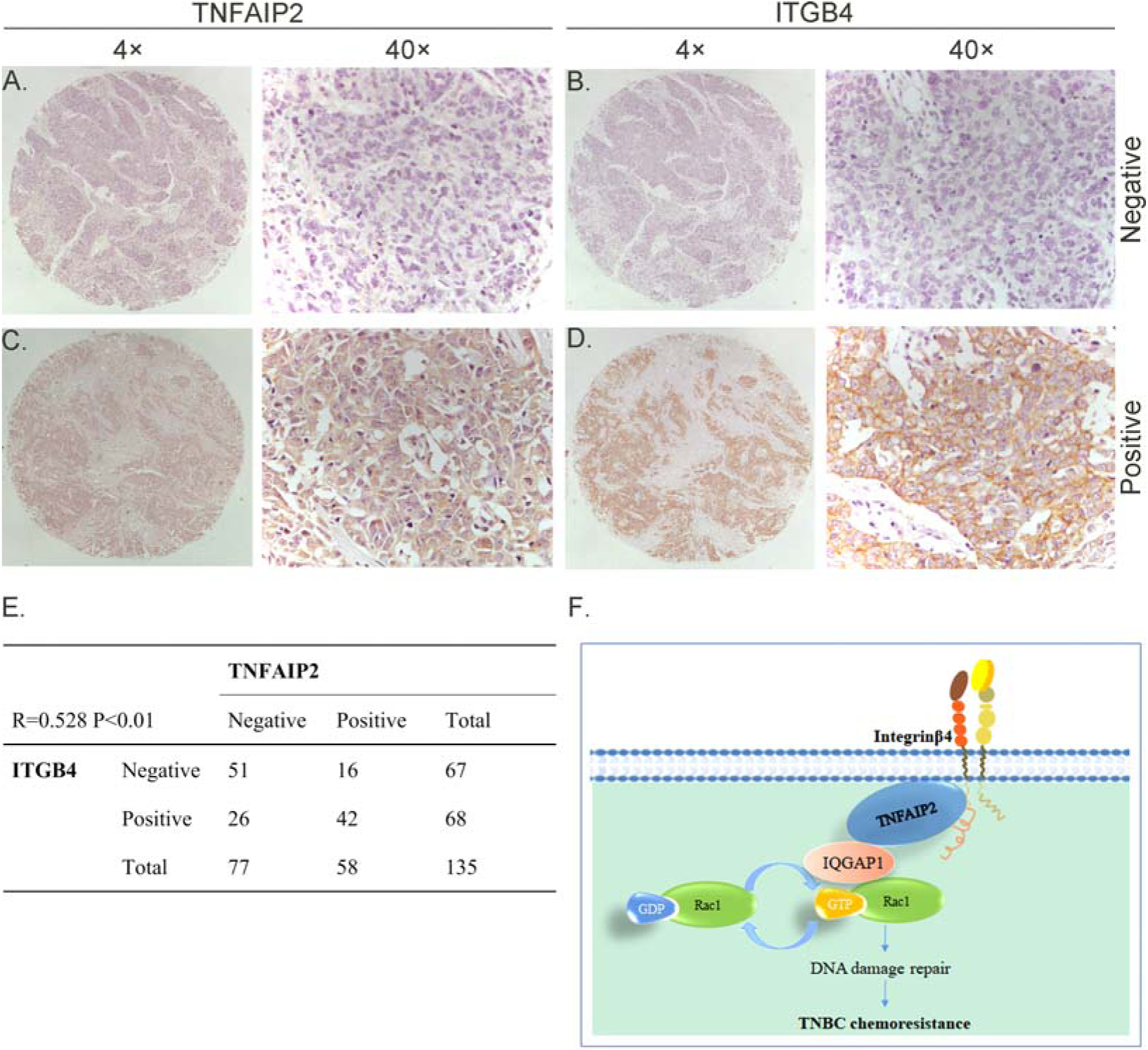
TNFAIP2 expression levels positively correlated with ITGB4 in TNBC tissues. Representative IHC images of TNFAIP2 and ITGB4 protein expression in breast cancer tissues are shown. The final expression assessment was performed by combining the two scores (0–2=low, 6–7=high). A and B indicate low scores, C and D indicate high scores, and E indicates that the TNFAIP2 and ITGB4 protein expression levels are positively correlated in human TNBC specimens. Figure F is the work model of this study.

## Discussion

Chemotherapies, including EPI and DDP, are the main choice for TNBC patients. Unfortunately, TNBC frequently develops resistance to chemotherapy[39]. Currently, PARP inhibitors are effective for TNBC with BRCA1/2 mutation or homologous recombination deficiency (HRD)[40–42]. PARP inhibitors can cause DNA damage repair defects and have synergistic lethal effects with HRD. Meanwhile, chemotherapy and PARP inhibitor resistance is also a major problem in the clinic.

In this study, we first found that TNFAIP2 promotes TNBC drug resistance and DNA damage repair through RAC1. Next, we found that TNFAIP2 interacts with IQGAP1and ITGB4. We verified that ITGB4 promotes TNBC drug resistance and DNA damage repair through the TNFAIP2/IQGAP1/RAC1 axis. Interestingly, we discovered for the first time that ITGB4 and TNFAIP2 promote RAC1 activity through IQGAP1. Our study reveals that ITGB4 promotes TNBC resistance through TNFAIP2-, IQGAP1-, and RAC1-mediated DNA damage repair (Figure 7). This study provides new targets for reversing TNBC resistance.

ITGB4 is well known to promote breast cancer stemness and can be activated by laminin 5[43]. In addition, ITGB4 is generally in partner with ITGA6, which is another marker of breast cancer stem cells[44] and drug resistance[43]. Therefore, whether ITGA6 has similar functions needs further study. It was reported that ITGB4 activates RAC1[45], but the mechanism is unclear. For the first time, we revealed that ITGB4 activates RAC1 through TNFAIP2 and IQGAP1. More importantly, ITGB4 promotes drug resistance through the TNFAIP2/IQGAP1/RAC1 axis.

TNFAIP2 plays important roles in different cellular and physiological processes, including cell proliferation, adhesion, migration, membrane TNT formation, angiogenesis, inflammation and tumorigenesis[14]. We previously found that TNFAIP2 was regulated by KLF5 and interacted with the small GTPases RAC1 and Cdc42, thereby regulating the actin cytoskeleton and cell morphology in breast cancer cells[9]. However, the detailed mechanism is not clearly understood. In this study, we found that IQGAP1 mediates this process. IQGAP1 is a crucial regulator of cancer development by scaffolding and facilitating different oncogenic pathways, especially RAC1/Cdc42, thus affecting proliferation, adhesion, migration, invasion, and metastasis[46]. In addition, IQGAP1 is increased during the differentiation of ovarian cancer stem cells and promotes aggressive behaviors[47]. In our study, we found that TNFAIP2 interacts with IQGAP1 and thus activates RAC1 to induce chemotherapy and PARP inhibitor drug resistance.

Furthermore, TNFAIP2 was reported to induce epithelial-to-mesenchymal transition and confer platinum resistance in urothelial cancer cells[12], and in embryonic stem cell (ESC) differentiation, TNFAIP2wasfound to be important in controlling lipid metabolism, which supports the ESC differentiation process and planarian organ maintenance[48]. Another study found that TNFAIP2 overexpression enhanced TNT-mediated autophagosome and lysosome exchange, preventing advanced glycation end product (AGE)-induced autophagy and lysosome dysfunction and apoptosis[49]. In cancer treatment, TNFAIP2 was chosen as one of the six signature genes predicting chemotherapeutic and immunotherapeutic efficacies, with high-senescore patients benefiting from immunotherapy and low-senescore patients responsive to chemotherapy[50].

These reports provide a possible explanation for previous studies showing that ITGB4 is important in EMT and cancer stemness. According to our results that there is an interaction between ITGB4 and TNFAIP2, ITGB4 might regulate EMT and stemness through TNFAIP2.TNFAIP2 is one of the important factors induced by tumor necrosis factor alpha (TNFα). Interestingly, TNFαrelease could be induced by therapeutic drugs from multiple tumor cell lines. The acquisition of docetaxel resistance was accompanied by increased constitutive production of TNFα[51]. In addition, TNFα is a key tumor-promoting effector molecule secreted by tumor-associated macrophages. *In vitro* neutralizing TNFα was observed to inhibit tumor progression and improve the curative effect of bevacizumab[52]. Therefore, the mechanism by which TNFα promotes chemotherapeutic resistance in breast cancer should be further investigated. For future studies, it will be important to develop *Tnfaip2* knockout mice to investigate the exact role of TNFAIP2 physiologically. According to recent studies and our findings, agents targeting the interaction among ITGB4/TNFAIP2/IQGAP1 would be a promising trend for developing drugs to overcome the resistance phenomenon.

In summary, ITGB4 and TNFAIP2 play important roles in breast cancer chemoresistance. TNFAIP2 activates RAC1 to promote chemoresistance through IQGAP1. In addition, ITGB4 activates RAC1 through TNFAIP2 and IQGAP1 and confer DNA damage-related drug resistance in TNBC (Figure 8F). These results indicate that the ITGB4/TNFAIP2/IQGAP1/RAC1 axis provides potential therapeutic targets to overcome DNA damage-related drug resistance in TNBC.

## Data availability statement

The authors confirm that the data supporting the findings of this study are available within the article and its supplementary materials.

## Acknowledgments

This work was supported by National Key R&D Program of China (2020YFA0112300), National Natural Science Foundation of China (81830087, U2102203, 81672624, 82102987, and 82203413), the Yunnan Fundamental Research Projects (202101AS070050), the Guangdong Foundation Committee for Basic and Applied Basic Research projects (2022A1515012420), and Yunnan (Kunming) Academician Expert Workstation (grant No. YSZJGZZ-2020025).

## Supplemental information titles and legends

**Figure 1-figure supplement 1.**
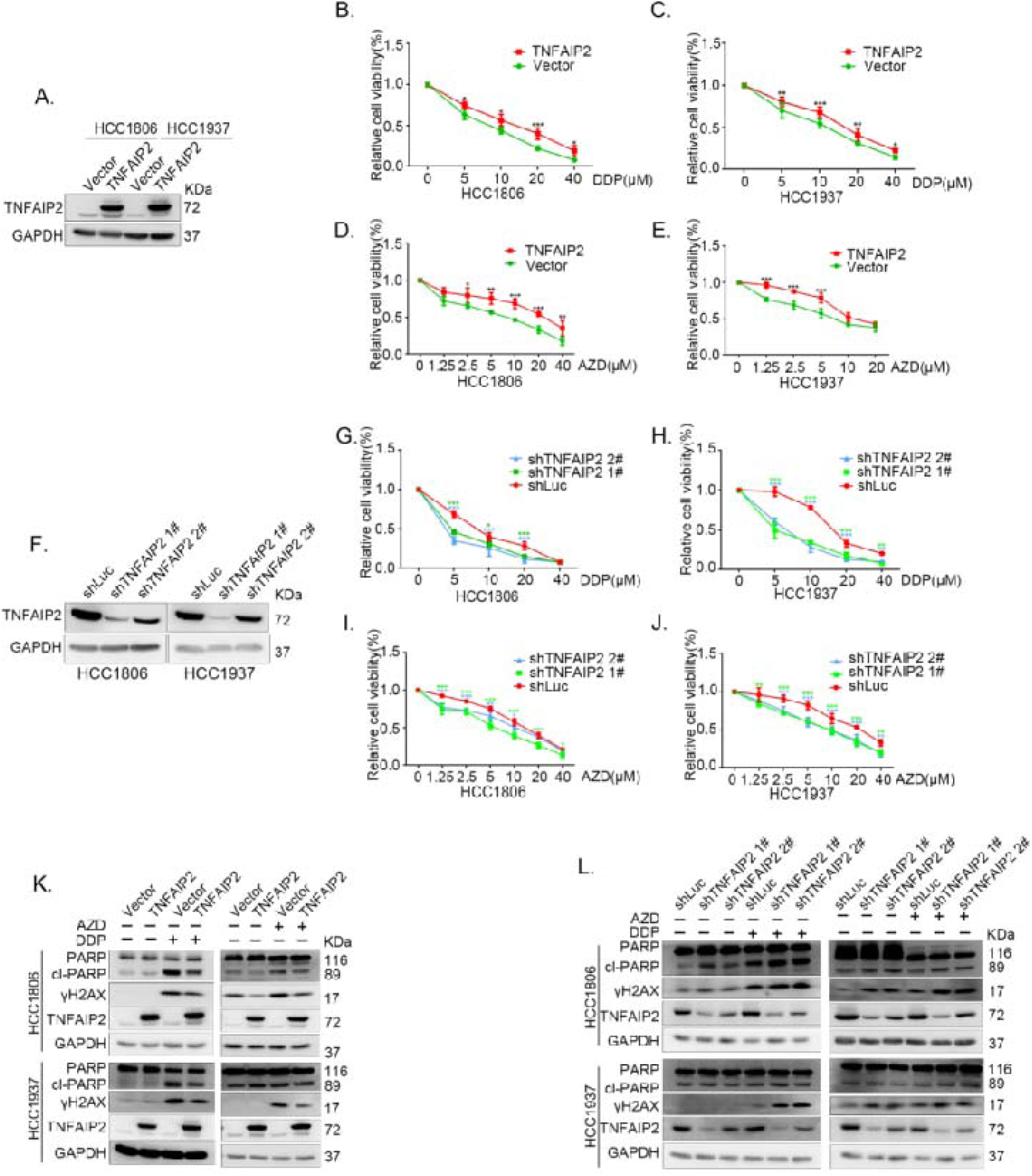
TNFAIP2 promotes TNBC DNA damage-related drug resistance. (A-E) Stable TNFAIP2 overexpression in HCC1806 and HCC1937 cells significantly increased cell viability in the presence of DDP (0-40μM) or AZD (0-40 μM) treatment for 48 h, as measured by the SRB assay. Statistical analysis was performed using one-way ANOVA, n=3-6, * *P*<0.05, ** *P*<0.01, *** *P*<0.001. TNFAIP2 protein expression was detected by WB. (F-J) Stable TNFAIP2 knockdown in HCC1806 and HCC1937 cells significantly decreased cell viability in the presence of DDP (0-40 μM) or AZD (0-40 μM) treatment for 48 h, as measured by the SRB assay. Statistical analysis was performed using one-way ANOVA, n=3, * *P*<0.05, ** *P*<0.01, *** *P*<0.001. TNFAIP2 protein expression was detected by WB. (K) TNFAIP2 promoted DNA damage repair in the presence of DDP and AZD. HCC1806 and HCC1937 cells stably overexpressing TNFAIP2 were treated with 20 μM DDP for 24 h or 48 h and 10 μM AZD for 24 h, respectively. TNFAIP2, γH2AX, and PARP protein expression was detected by WB. (L) TNFAIP2 knockdown increased DNA damage in the presence of DDP and AZD. Stable TNFAIP2 knockdown cells were treated with 2.5 or 20 μM DDP for 24 h and 2.5 μM AZD for 24 h. TNFAIP2, γH2AX, and PARP protein expression was detected by WB. Figure 1-figure supplement 1-source data 1 Uncropped western blot images for Figure 1-figure supplement 1

**Figure 2-figure supplement 1.**
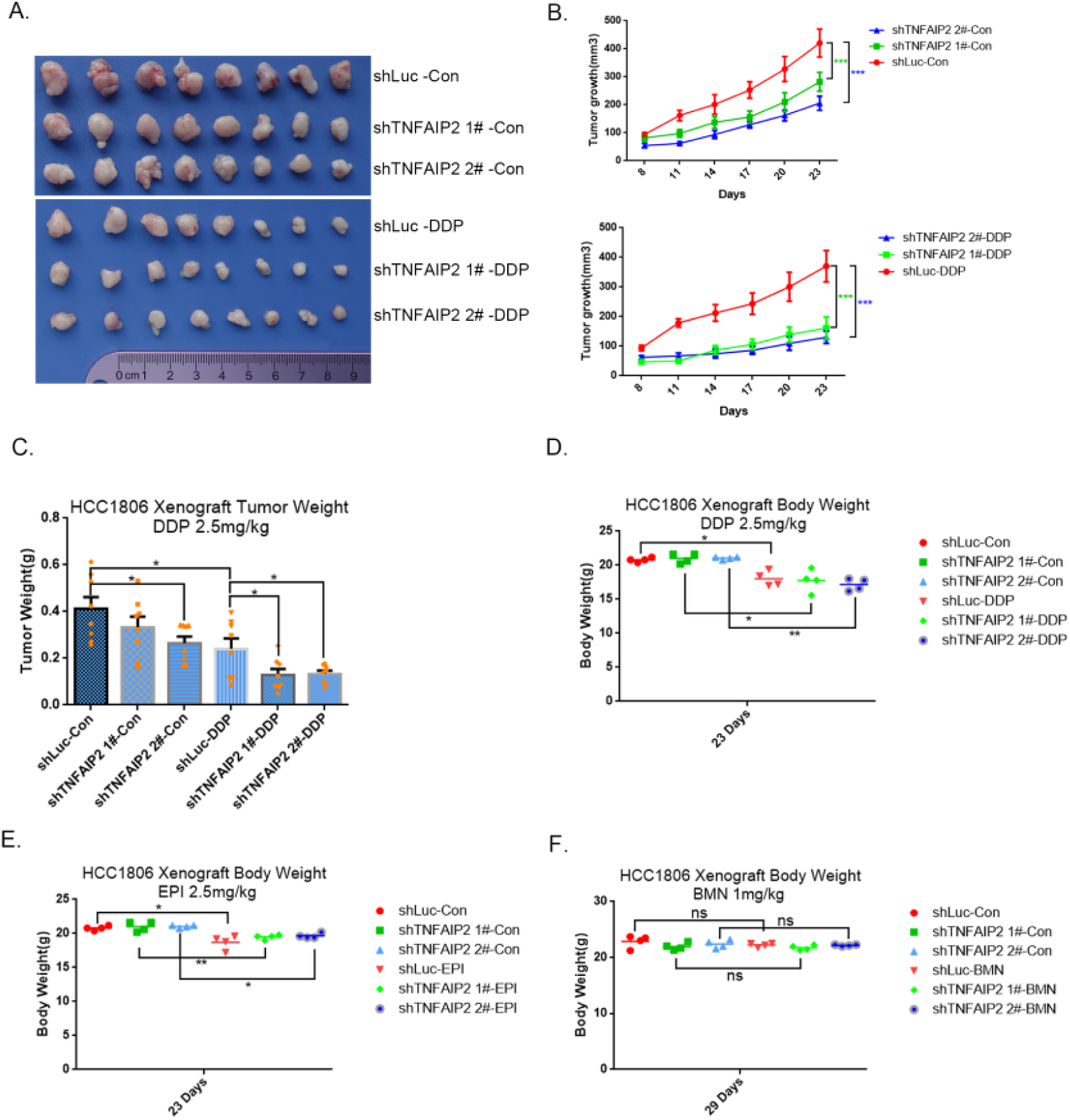
TNFAIP2 confers TNBC drug resistance *in vivo*. (A-F) TNFAIP2 knockdown increased the sensitivity ofHCC1806 breast cancer cells to DDP *in vivo*. HCC1806 cells with stable TNFAIP2 knockdown were transplanted into the fat pad of 7-week-old female nude mice. When the average tumor size reached approximately 50 mm^3^ after inoculation, mice in each group were randomly divided into two subgroups (n=4/group) to receive DDP (2.5 mg/kg) or vehicle control for 23 days (A). Tumor size was measured twice a week (B), tumor masses were collected and weighed at the end of the experiments (C), and mouse masses were collected and weighed at the beginning or end of the experiments (D-F). *: *P*<0.05, **: *P*<0.01, ***: *P*<0.001, *t-*test.

**Figure 2-figure supplement 2.**
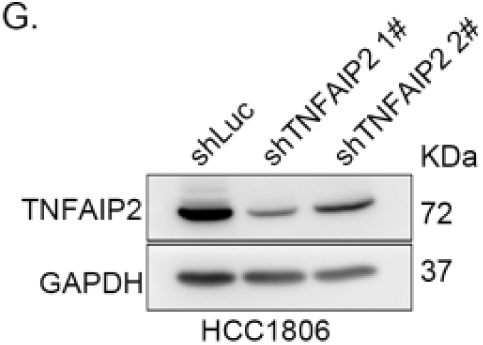
TNFAIP2 confers TNBC drug resistance *in vivo*. (G) TNFAIP2 was stably knocked down in HCC1806, as determined by Western blotting. Figure 2-figure supplement 2-source data 1 Uncropped western blot images for Figure 2-figure supplement 2.

**Figure 3-figure supplement 1.**
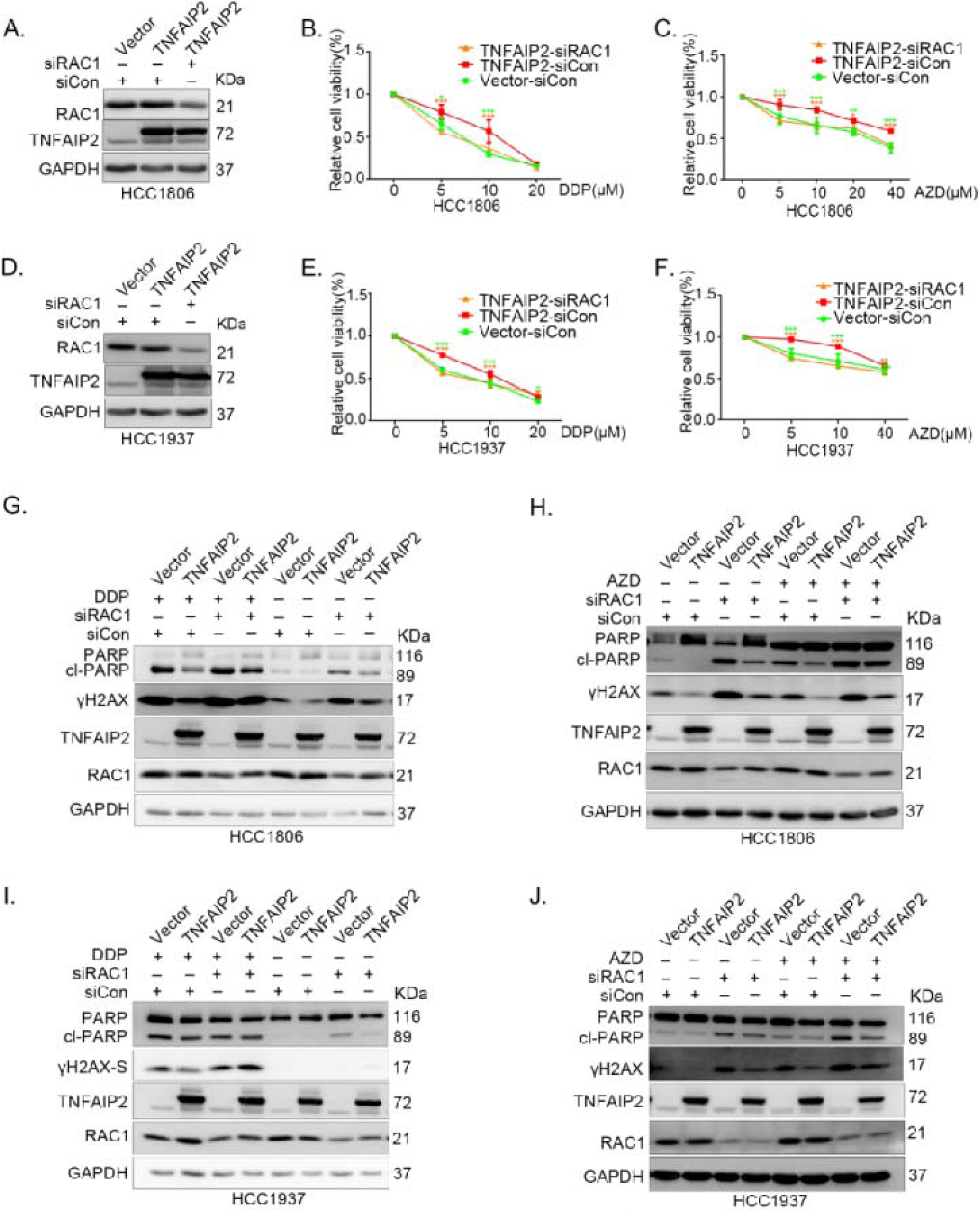
TNFAIP2 promotes TNBC drug resistance and DNA damage repair via RAC1. (A-F) RAC1 knockdown abolished TNFAIP2-induced TNBC resistance to DDP and AZD. HCC1806 (A-C) and HCC1937 (D-F) cells with stable TNFAIP2 overexpression were transfected with RAC1 or control siRNA, followed by treatment with DDP (0-40 μM) and AZD (0-40 μM) for 48 or 72 h, respectively. Cell viability was measured by the SRB assay. Statistical analysis was performed using one-way ANOVA, n=3-4, * *P*<0.05, ** **P**<0.01, *** *P*<0.001. Protein expression levels were analyzed by WB. (G-J) RAC1 depletion abolished TNFAIP2-induced DNA damage decrease in response to DDP and AZD. HCC1806 (G-H) and HCC1937 (I-J) cells with stable TNFAIP2 overexpression were transfected with RAC1 or control siRNA, followed by treatment with DDP (20 μM) and AZD (10 μM) for 24 h, respectively. Protein expression levels were analyzed by WB. Figure 3-figure supplement 1-source data 1 Uncropped western blot images for Figure 3-figure supplement 1

**Figure 4-figure supplement 1.**
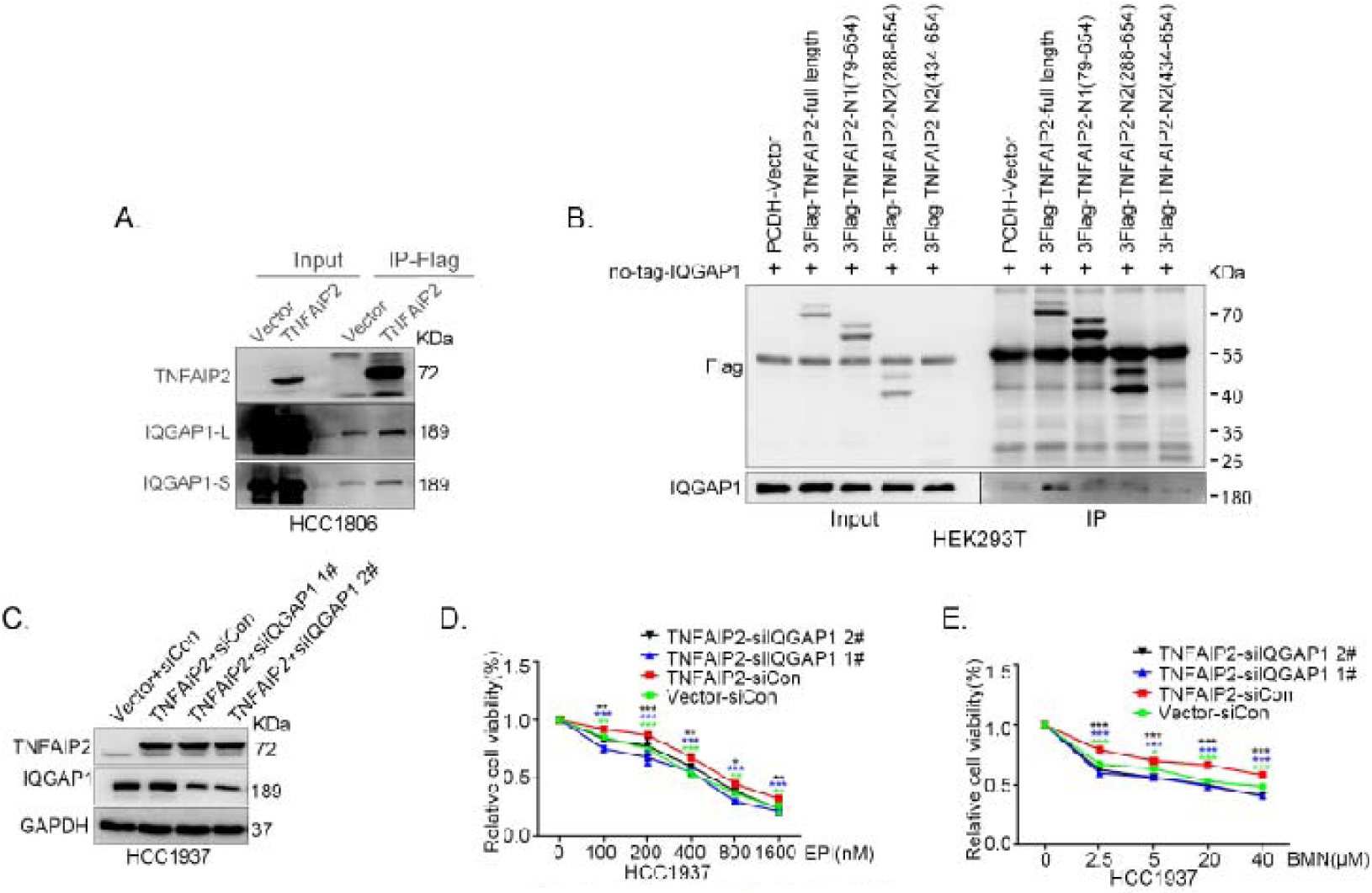
IQGAP1 mediates RAC1 activation by TNFAIP2 and promotes TNBC drug resistance. (A) TNFAIP2 interacts with IQGAP1.HCC1806 cells with stable TNFAIP2 overexpression were collected from flag-tagged TNFAIP2 cell lysates for immunoprecipitation assays using Flag-M2 beads, and IQGAP1 was detected by WB. (B) Mapping the domains of TNFAIP2 that interact with IQGAP1. Flag-tagged full-length or truncated TNFAIP2 was transfected into HEK293T cells with no-tagged full-length IQGAP1. Cell lysates were collected for immunoprecipitation using Flag-M2 beads, and IQGAP1 was detected by WB. (C-E) IQGAP1 knockdown abolished TNFAIP2-confered resistance to EPI and BMN. HCC1937 cells with stable TNFAIP2 overexpression were transfected with IQGAP1 or control siRNA, followed by treatment with EPI (0-1600 nM) and BMN (0-40 μM) for 48 or 72 h, respectively. Cell viability was measured by the SRB assay. Statistical analysis was performed using one-way ANOVA, n=3, * *P*<0.05, ** *P*<0.01, *** *P*<0.001. Protein expression levels were analyzed by WB. Figure 4-figure supplement 1-source data 1 Uncropped western blot images for Figure 4-figure supplement 1

**Figure 5-figure supplement 1.**
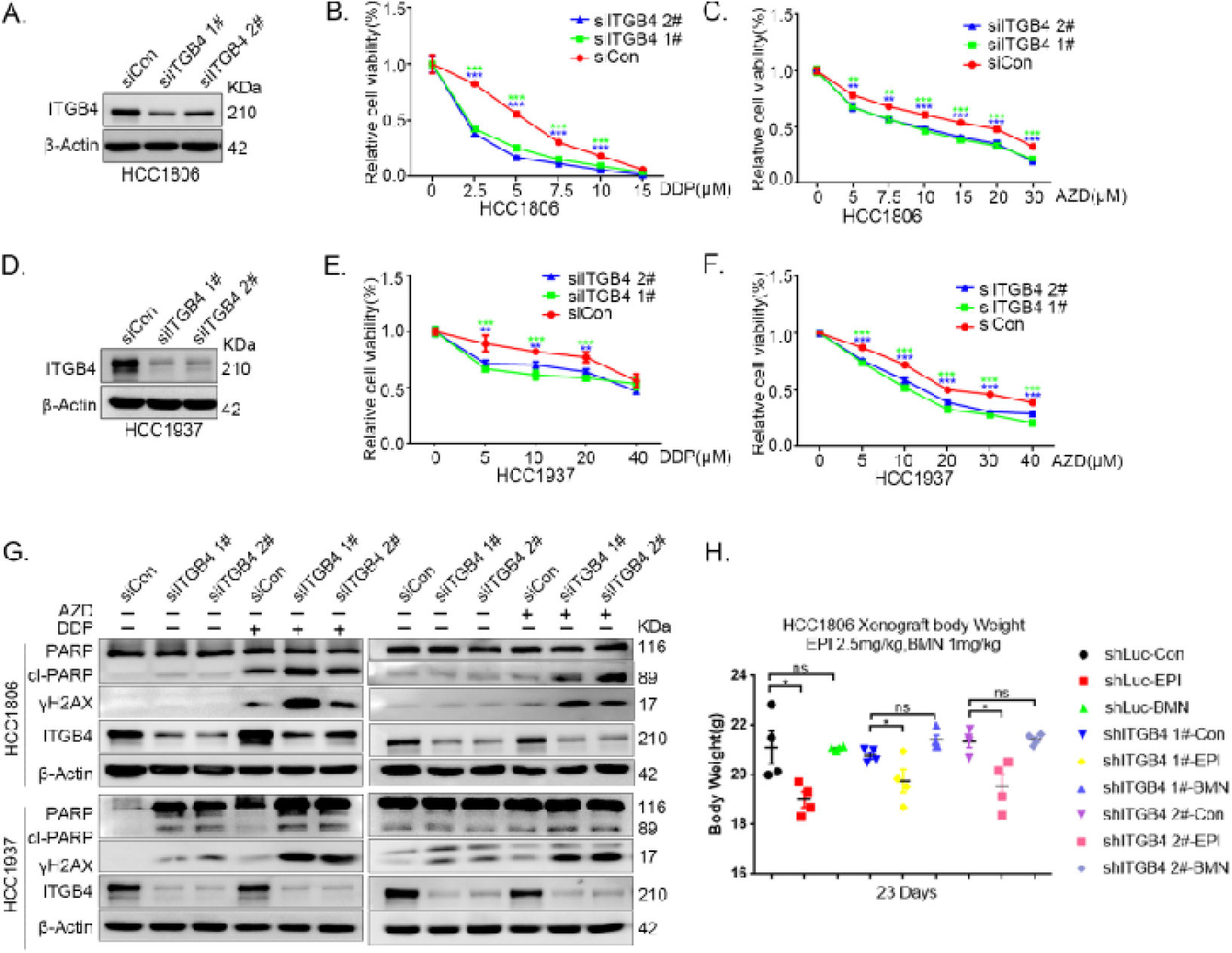
ITGB4 interacts with TNFAIP2 and promotes TNBC drug resistance and DNA damage repair. (A-F) ITGB4 knockdown in HCC1806 and HCC1937 cells significantly decreased cell viability in the presence of DDP (0-40 μM) and AZD (0-40 μM), as measured by the SRB assay. Statistical analysis was performed using one-way ANOVA, n=3, * *P*<0.05, ** *P*<0.01, *** *P*<0.001. ITGB4 protein expression was detected by WB. (G) ITGB4 knockdown increased DNA damage of DDP and AZD. HCC1806 and HCC1937 cells with ITGB4 knockdown were treated with 5 μM or 7.5 μM DDP for 24 h and 15 μM or 20 μM AZD for 24 h, respectively. ITGB4, γH2AX, and PARP protein expression was detected by WB. (H) Mouse masses were collected and weighed at the end of the experiments. Figure 5-figure supplement 1-source data 1 Uncropped western blot images for Figure 5-figure supplement 1

**Figure 5-figure supplement 2.**
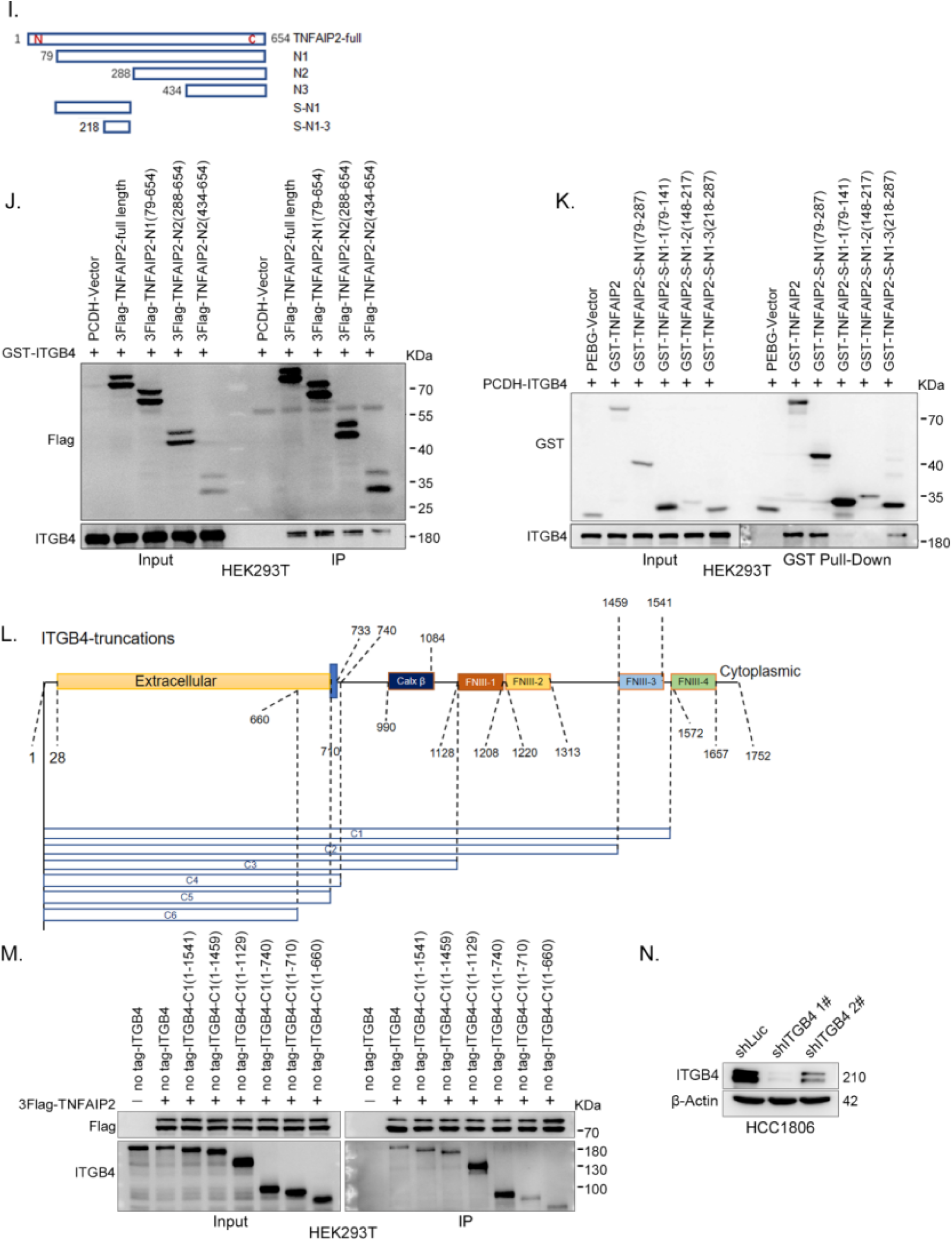
ITGB4 interacts with TNFAIP2 and promotes TNBC drug resistance and DNA damage repair. (I) The model of full-length or truncated TNFAIP2. (J) Mapping the domains of TNFAIP2 that interact with ITGB4. GST-tagged Full-length ITGB4 was transfected into HEK293T cells with flag-tagged full-length or truncated TNFAIP2. TNFAIP2 protein was immunoprecipitated using Flag-M2 beads, and ITGB4 was detected by WB. (K) Mapping the domains of TNFAIP2 that interact with ITGB4. No-tagged full-length ITGB4 was transfected into HEK293T cells with GST-tagged full-length or truncated TNFAIP2. Cell lysates were collected for the GST pull-down assay, and ITGB4 was detected by WB. (L) The model of full-length or truncated ITGB4. (M) Mapping the domains of ITGB4 that interact with TNFAIP2. Flag-tagged full-length TNFAIP2 was transfected into HEK293T cells with no-tagged full-length or truncated ITGB4. Cell lysates were collected for immunoprecipitation using Flag-M2 beads, and ITGB4 was detected by WB. (N) ITGB4 was stably knocked down in HCC1806, as determined by Western blotting. Figure 5-figure supplement 2-source data 1 Uncropped western blot images for Figure 5-figure supplement 2

**Figure 7-figure supplement 1.**
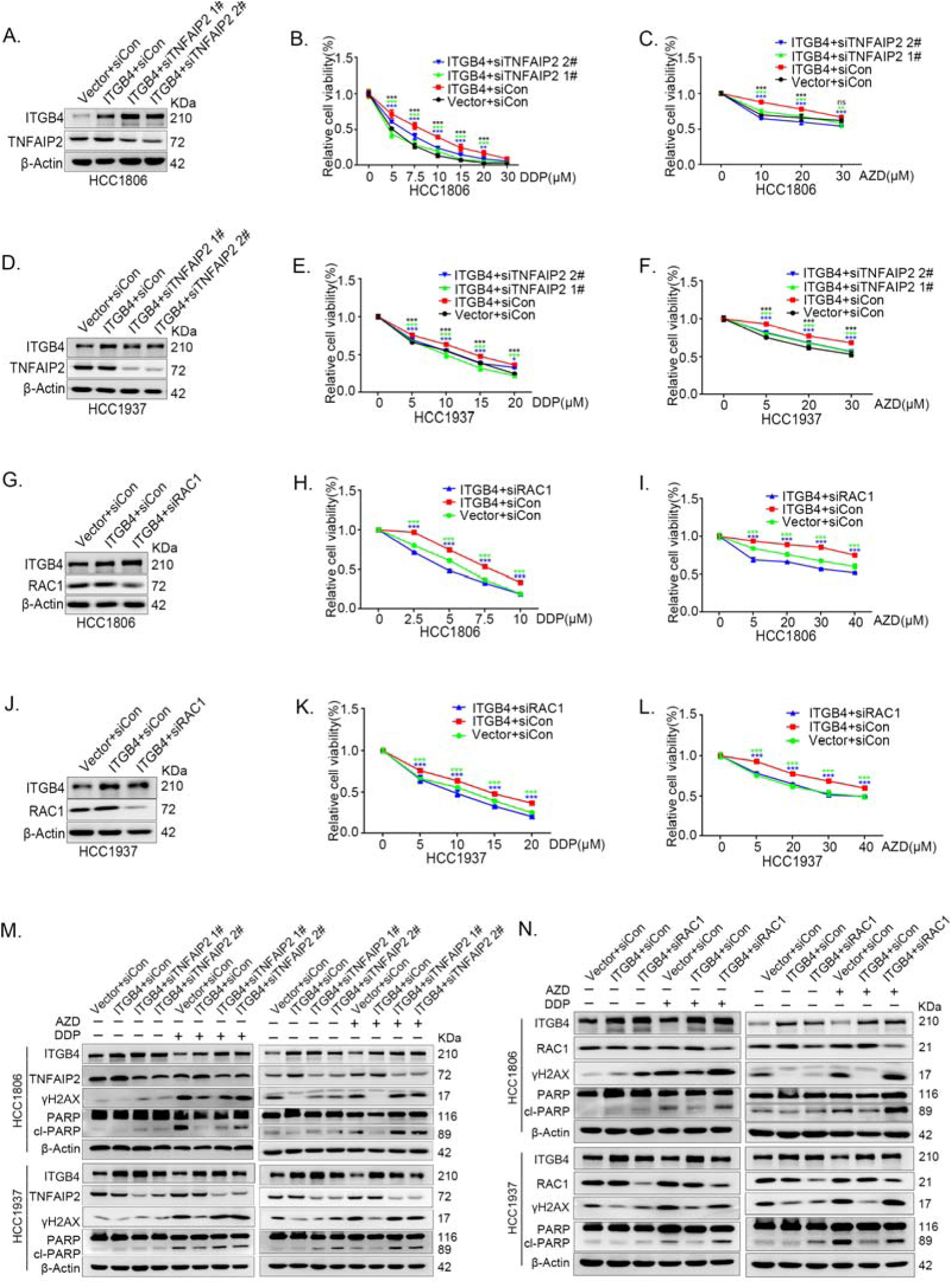
ITGB4 promotes TNBC drug resistance via TNFAIP2/IQGAP1/RAC1. (A-F) ITGB4 promotes TNBC drug resistance through TNFAIP2. TNFAIP2 knockdown abolished ITGB4-induced resistance to DDP and AZD. HCC1806 (A-C) and HCC1937 (D-F) cells with stable ITGB4 overexpression were transfected with TNFAIP2 or control siRNA, followed by treatment with DDP (0-30 μM) and AZD (0-30 μM) for 48 or 72 h, respectively. Cell viability was measured by the SRB assay. Protein expression levels were analyzed by WB. (G-L) ITGB4 promotes TNBC drug resistance through RAC1. RAC1 knockdown abolished ITGB4-induced resistance to DDP and AZD. HCC1806 (G-I) and HCC1937 (J-L) cells with stable ITGB4 overexpression were transfected with RAC1 or control siRNA, followed by treatment with DDP (0-20 μM) and AZD (0-40 μM) for 48 or 72 h, respectively. Cell viability was measured by the SRB assay. Statistical analysis was performed using one-way ANOVA, n=3, * *P*<0.05, ** *P*<0.01, *** *P*<0.001. Protein expression levels were analyzed by WB. (M) ITGB4 promotes DNA damage repair through TNFAIP2. HCC1806 and HCC1937 cells with stable ITGB4 overexpression were transfected with TNFAIP2 or control siRNA, followed by treatment with DDP (7.5 μM or 10 μM) and AZD (20 μM or 30 μM) for 24 h. Protein expression levels were analyzed by WB. (N) ITGB4 promotes DNA damage repair through RAC1. HCC1806 and HCC1937 cells with stable ITGB4 overexpression were transfected with RAC1 or control siRNA, followed by treatment with DDP (7.5 μM) and AZD (20 μM or 30 μM) for 24 h. Protein expression levels were analyzed by WB. Figure 7-figure supplement 1-source data 1 Uncropped western blot images for Figure 7-figure supplement 1

**Figure 7-figure supplement 2.**
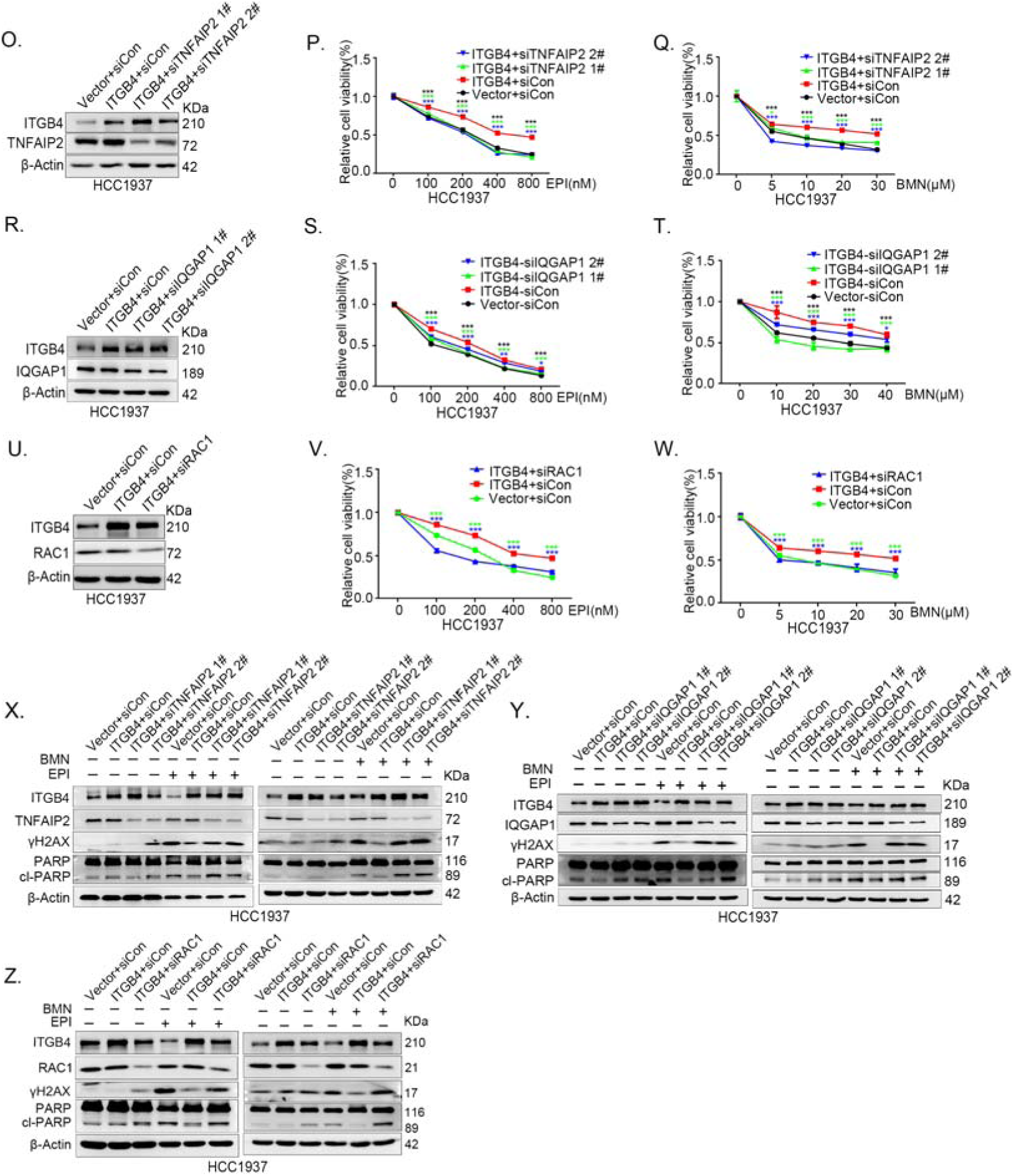
ITGB4 promotes TNBC drug resistance via TNFAIP2/IQGAP1/RAC1. (O-Q) ITGB4 promotes TNBC drug resistance through TNFAIP2. TNFAIP2 knockdown abolished ITGB4-induced resistance to EPI and BMN. HCC1937 cells with stable ITGB4 overexpression were transfected with TNFAIP2 or control siRNA, followed by treatment with EPI (0-800 nM) and BMN (0-30 μM) for 48 or 72 h, respectively. Cell viability was measured by the SRB assay. Statistical analysis was performed using one-way ANOVA, n=3, * *P*<0.05, ** *P*<0.01, *** *P*<0.001. Protein expression levels were analyzed by WB. (R-T) ITGB4 promotes TNBC drug resistance through IQGAP1. HCC1937 cells with stable ITGB4 overexpression were transfected with IQGAP1 or control siRNA, followed by treatment with EPI (0-800 nM) and BMN (0-40 μM) for 48 or 72 h, respectively. Cell viability was measured by the SRB assay. Statistical analysis was performed using one-way ANOVA, n=3, * *P*<0.05, ** *P*<0.01, *** *P*<0.001. Protein expression levels were analyzed by WB. (U-W) ITGB4 promotes TNBC drug resistance through RAC1. HCC1937 cells with stable ITGB4 overexpression were transfected with RAC1 or control siRNA, followed by treatment with EPI (0-800 nM) and BMN (0-30 μM) for 48 or 72 h, respectively. Cell viability was measured by the SRB assay. Statistical analysis was performed using one-way ANOVA, n=3, * *P*<0.05, ** *P*<0.01, *** *P*<0.001. Protein expression levels were analyzed by WB. (X) ITGB4 promotes DNA damage repair through TNFAIP2. HCC1937 cells with stable ITGB4 overexpression were transfected with TNFAIP2 or control siRNA, followed by treatment with EPI (800 nM) and BMN (10 μM) for 24h. Protein expression levels were analyzed by WB. (Y) ITGB4 promotes DNA damage repair through IQGAP1. HCC1937 cells with stable ITGB4 overexpression were transfected with IQGAP1 or control siRNA, followed by treatment with EPI (800 nM) and BMN (10 μM) for 24 h. Protein expression levels were analyzed by WB. (Z) ITGB4 promotes DNA damage repair through RAC1. HCC1937 cells with stable ITGB4 overexpression were transfected with RAC1 or control siRNA, followed by treatment with EPI (800 nM) and BMN (10 μM) for 24 h. Protein expression levels were analyzed by WB. Figure 7-figure supplement 2-source data 1 Uncropped western blot images for Figure 7-figure supplement 2

## Materials and Methods

### Cell lines and reagents

All cell lines used in this study, including HCC1806, HCC1937, and HEK293T cells, were purchased from ATCC (American Type Culture Collection, Manassas, VA, USA) and validated by STR (short tandem repeat) analysis (supplementary file 1) and these cell lines tested negative for mycoplasma contamination (supplementary file 2). HCC1806 and HCC1937 cells were cultured in RPMI 1640 medium supplemented with 5% FBS. HEK293T cells were cultured in DMEM (Thermo Fisher, Grand Island, USA) with 5% FBS at 37°Cwith 5% CO_2_.Epirubicin(EPI) (Cat#HY-13624A), cisplatin(DDP)(Cat#HY-17394), talazoparib(BMN) (Cat#HY-16106),and olaparib(AZD) (Cat#HY-10162) were purchased from MCE (New Jersey, USA).

### Plasmid construction and stable TNFAIP2 and ITGB4 overexpression

We constructed the full-length *TNFAIP2/ITGB4* gene and then subcloned them into the pCDH lentiviral vector. The packaging plasmids (including pMDLg/pRRE, pRSV-Rev, and pCMV-VSV-G) and pCDH-TNFAIP2/ITGB4 expressionplasmid were cotransfected into HEK293T cells (2 ×10^6^ in 10 cm plate) to produce lentivirus. Following transfection for 48 h, the lentivirus was collected and used to infect HCC1806 and HCC1937 cells. Forty-eight hours later, puromycin (2 μg/ml) was used to screen the cell populations.

### Stable knockdown of TNFAIP2 and ITGB4

The pSIH1-H1-puro shRNA vector was used to express TNFAIP2, ITGB4 and luciferase(LUC)shRNAs.*TNFAIP2*shRNA#1,5’-GACUUGGGCUCACAGAUAA-3’;*TNFAIP2*shRNA#2,5’-GAUUGAGGUGGCCACUUAU-3’;*ITGB4*shRNA#1,5’-ACGACAGCTTCCTTATGTA-3’;*ITGB4*shRNA#2,5’-CAGCGACTACACTATTG GA-3’;*Luciferase*shRNA,5’-CUUACGCUGAGUACUUCGA-3’; HCC1806 and HCC1937 cells were infected with lentivirus. Stable populations were selected using 1 to 2 mg/mL puromycin. The knockdown effect was evaluated by Western blotting.

### RNA interference

The siRNA target sequences used in this study are as follows:*TNFAIP2*siRNA#1,5’-GACUUGGGCUCACAGAUAA-3’;*TNFAIP2*siRN A#2,5’-GAUUGAGGUGGCCACUUAU-3’;*ITGB4*siRNA#1,5’-ACGACAGCTTC CTTATGTA-3’;*ITGB4*siRNA#2,5’-CAGCGACTACACTATTGGA-3’;*RAC1*siRNA,5’-CGGCACCACUGUCCCAACA-3’;*IQGAP1*siRNA#1,5’-GCAGGTGGATTAC TATAAA-3’;*IQGAP1*siRNA#2,5’-CUAGUGAAACUGCAACAGA-3’. All siRNAs were synthesized by RiboBio (RiboBio, China) and transfected at a final concentration of 50 nM.

### Antibodies and Western blotting (WB)

The WB procedure has been described in our previous study[53]. Anti-TNFAIP2 (sc-28318), anti-ITGB4 (sc-9090) and anti-GAPDH (sc-25778) antibodies were purchased from Santa Cruz Biotechnology (Santa Cruz, CA, USA). The anti-PARP (#9542) antibody was purchased from CST. Anti-RAC1 (05–389) and anti-γH2AX (3475627)antibodies were purchased from Millipore (Billerica, MA,USA). Anti-β-actin (A5441) and anti-Tubulin (T5168) antibodies were purchased from Sigma[Aldrich (St Louis, MO, USA). The anti-IQGAP1 (ab86064) antibody was purchased from Abcam.

### Immunoprecipitation and silver staining

Immunoprecipitation and silver staining lysates from HCC1806 cells stably expressing Flag-TNFAIP2 were prepared by incubating the cells in lysis buffer containing a protease inhibitor cocktail (MCE). Cell lysates were obtained from approximately 2.5×10^8^ cells, and after binding with anti-Flag M2 affinity gel (Sigma) for 2 h as recommended by the manufacturer, the affinity gel was washed with cold lysis plus 0.2% NP-40. FLAG peptide (Sigma) was applied to elute the Flag-labeled protein complex as described by the vendor. The elutes were collected and visualized on NuPAGE 4%-12% Bis-Trisgels (Invitrogen, CA, USA) followed by silver staining with a silver staining kit (Pierce, Illinois, USA). The distinct protein bands were retrieved and analyzed by LC Mass (supplementary table 1).

### Immunoprecipitation and GST pull-down

For exogenous interaction between ITGB4 and Flag-TNFAIP2, cell lysates were directly incubated with anti-Flag M2 affinity gel (A2220; Sigm) overnight at 4°C. For endogenous protein interaction, cell lysates were first incubated with anti-TNFAIP2/ITGB4/IQGAP1 antibodies or mouse IgG/rabbit IgG (sc-2028;Santa Cruz Biotechnology, CA, USA) and then incubated with Protein A/G plusagarose beads (sc-2003; Santa Cruz Biotechnology For the GST pull down assay, cell lysates were directly incubated with GlutathioneSepharose 4B (52-2303-00; GE Healthcare) overnight at 4°C. The precipitates were washed four times with 1 ml of lysis buffer, boiled for 10 minutes with 1×SDS sample buffer, and subjected to WB analysis.

### Cell viability assays

Cell viability was measured by SRB assays as described in our previous study[54]. Cell viability was measured by SRB assays. Briefly, cells were seeded in 96-well plates. Then, the cells were cultured for the indicated time and fixed with 10% trichloroacetic acid at room temperature for30 min, followed by incubation with 0.4% SRB (w/v) solution in 1% acetic acid for 20 min at room temperature. Finally, SRB was dissolved in 10 mM unbuffered Tris base, and the absorbance was measured at a wavelength of 530 nm on a plate reader (Bio Tek, Vermont, USA).

### RAC1 activation assays

RAC1 activation was examined using the Cdc42 Activation Assay Biochem Kit (BK034, Cytoskeleton, Denver, USA) following them anufacturer’s instructions. Cells were harvested with cell lysis buffer, and1 mg of protein lysate in a 1 ml total volume at 4°C was immediately precipitated with 10 μg of PAK-PBD beads for 60 min with rotation. After washing three times with wash buffer, agarose beads were resuspended in 30 μl of 2×SDS sample buffer and boiled for 5 min. RAC1-GTP was examined by WB with an anti-RAC1 antibody.

### Xenograft experiments

We purchased 6- to 7-week-old female BALB/cnude mice from SLACCAS (Changsha, China). HCC1806-shLuc, HCC1806-shTNFAIP2, or HCC1806-shITGB4 cells (1 × 10^6^ in Matrigel (BD Biosciences, NY, USA)) were implanted into the mammary fat pads of the mice. When the tumor volume reached approximately 50 mm^3^, the nude mice were randomly assigned to the control and treatment groups (n = 4/group). EPI, BMN, and DDP were dissolved in ddH_2_O. The control group was given vehicle alone, and the treatment group received EPI (2.5 mg/kg), BMN (1 mg/kg), and DDP (2.5 mg/kg) alone via intraperitoneal injection every three days for 18 or 27 days. The tumor volume was calculated as follows: tumor volume was calculated by the formula: (π×length × width^2^)/6.

### Immunohistochemical staining

Paraffin-embedded clinical TNBC specimens were obtained from the First Affiliated Hospital, Zhengzhou University, Zhengzhou, China. Informed consent was obtained from all subjects. Two tissue microarrays containing 135 TNBC breast cancer tissues were constructed. For the immunohistochemistry (IHC) assay, the slides were deparaffinized, rehydrated, and pressure cooker heated for 2.5 min in EDTA for antigen retrieval. Endogenous peroxidase activity was inactivated by adding an endogenous peroxidase blocker (OriGene, China) for 15 min at room temperature. Slides were incubated overnight at 4°C with anti-TNFAIP2 (1:200) or anti-ITGB4 (1:500). After 12 h, the slides were washed three times with PBS and incubated with secondary antibodies (hypersensitive enzyme-labeled goat anti-mouse/rabbit IgG polymer (OriGene, China) at room temperature for 20 min, DAB concentrate chromogenic solution (1:200dilution of concentrated DAB chromogenic solution), counterstained with 0.5% hematoxylin, dehydrated with graded concentrations of ethanol for 3 min each (70%–80%–90%–100%),and finally stained with dimethyl benzene immunostainedslides were evaluated by light microscopy. The IHC signal was scored using the ‘Allred Score’ method.

### Statistical analysis

All graphs were created using GraphPad Prism software version 8.0. Comparisons between two independent groups were assessed by two-tailed Student’s *t*-test. One-way analysis of variance with least significant differences was used for multiple group comparisons. *P-*values of <0.05, 0.01 or 0.001 were considered to indicate a statistically significant result, comparisons significant at the 0.05 level are indicated by*, at the 0.01 level are indicated by **, or at the 0.001 level are indicated by ***.

### Ethics

Animal feeding and experiments were approved by the animal ethics committee of the affiliated Hospital of Guangdong Medical university (GDY2102096, supplementary file 3). Clinical samples were approved by the relevant institution (YS2021036, supplementary file 4).

## References

1. Bray F, Ferlay J, Soerjomataram I, Siegel RL, Torre LA, Jemal A. Global cancer statistics 2018: GLOBOCAN estimates of incidence and mortality worldwide for 36 cancers in 185 countries. CA Cancer J Clin. 2018;68: 394–424.

2. Zhao S, Ma D, Xiao Y, et al. Molecular Subtyping of Triple-Negative Breast Cancers by Immunohistochemistry: Molecular Basis and Clinical Relevance 2020;25: e1481–e1491.

3. Sotiriou C, Pusztai L. Gene-expression signatures in breast cancer. N Engl J Med. 2009;360: 790–800.

4. Bai X, Ni J, Beretov J, Graham P, Li Y. Triple-negative breast cancer therapeutic resistance: Where is the Achilles’ heel? Cancer Lett. 2021;497: 100–111.

5. Jamdade VS, Sethi N, Mundhe NA, Kumar P, Lahkar M, Sinha N. Therapeutic targets of triple-negative breast cancer: a review. Br J Pharmacol. 2015;172: 4228–4237.

6. Hill DP, Harper A, Malcolm J, et al. Cisplatin-resistant triple-negative breast cancer subtypes: multiple mechanisms of resistance. BMC Cancer. 2019;19: 1039.

7. Longley DB, Johnston PG. Molecular mechanisms of drug resistance. J Pathol. 2005;205: 275–292.

8. Rincón R, Zazo S, Chamizo C, et al. c-Jun N-Terminal Kinase Inactivation by Mitogen-Activated Protein Kinase Phosphatase 1 Determines Resistance to Taxanes and Anthracyclines in Breast Cancer. Mol Cancer Ther. 2016;15: 2780–2790.

9. Jia L, Zhou Z, Liang H, et al. KLF5 promotes breast cancer proliferation, migration and invasion in part by upregulating the transcription of TNFAIP2. Oncogene. 2016;35: 2040–2051.

10. Chen LC, Chen CC, Liang Y, Tsang NM, Chang YS, Hsueh C. A novel role for TNFAIP2: its correlation with invasion and metastasis in nasopharyngeal carcinoma. Mod Pathol. 2011;24: 175–184.

11. Cheng Z, Wang HZ, Li X, et al. MicroRNA-184 inhibits cell proliferation and invasion, and specifically targets TNFAIP2 in Glioma. J Exp Clin Cancer Res. 2015;34: 27.

12. Niwa N, Tanaka N. TNFAIP2 expression induces epithelial-to-mesenchymal transition and confers platinum resistance in urothelial cancer cells 2019;99: 1702–1713.

13. Xie Y, Wang B. Downregulation of TNFAIP2 suppresses proliferation and metastasis in esophageal squamous cell carcinoma through activation of the Wnt/β-catenin signaling pathway. Oncol Rep. 2017;37: 2920–2928.

14. Jia L, Shi Y, Wen Y, Li W, Feng J, Chen C. The roles of TNFAIP2 in cancers and infectious diseases 2018;22: 5188–5195.

15. Didsbury J, Weber RF, Bokoch GM, Evans T, Snyderman R. rac, a novel ras-related family of proteins that are botulinum toxin substrates. J Biol Chem. 1989;264: 16378–16382.

16. Rul W, Zugasti O, Roux P, et al. Activation of ERK, controlled by Rac1 and Cdc42 via Akt, is required for anoikis. Ann N Y Acad Sci. 2002;973: 145–148.

17. Liu W, Han J, Shi S, Dai Y, He J. TUFT1 promotes metastasis and chemoresistance in triple negative breast cancer through the TUFT1/Rab5/Rac1 pathway. Cancer Cell Int. 2019;19: 242.

18. Yan Y, Greer PM, Cao PT, Kolb RH, Cowan KH. RAC1 GTPase plays an important role in γ-irradiation induced G2/M checkpoint activation. Breast Cancer Res. 2012;14: R60.

19. Wu M, Li L, Hamaker M, Small D, Duffield AS. FLT3-ITD cooperates with Rac1 to modulate the sensitivity of leukemic cells to chemotherapeutic agents via regulation of DNA repair pathways. Haematologica. 2019;104: 2418–2428.

20. Hu H, Juvekar A, Lyssiotis CA, et al. Phosphoinositide 3-Kinase Regulates Glycolysis through Mobilization of Aldolase from the Actin Cytoskeleton. Cell. 2016;164: 433–446.

21. Li Q, Qin T, Bi Z, et al. Rac1 activates non-oxidative pentose phosphate pathway to induce chemoresistance of breast cancer 2020;11: 1456.

22. Hervieu A, Heuss SF. A PI3K- and GTPase-independent Rac1-mTOR mechanism mediates MET-driven anchorage-independent cell growth but not migration 2020;13.

23. Li Y, Li P, Wang N. Effect of let-7c on the PI3K/Akt/FoxO signaling pathway in hepatocellular carcinoma. Oncol Lett. 2021;21: 96.

24. Feng YX, Zhao JS, Li JJ, et al. Liver cancer: EphrinA2 promotes tumorigenicity through Rac1/Akt/NF-kappaB signaling pathway. Hepatology. 2010;51: 535–544.

25. Higuchi M, Onishi K, Kikuchi C, Gotoh Y. Scaffolding function of PAK in the PDK1-Akt pathway. Nat Cell Biol. 2008;10: 1356–1364.

26. Hu F, Li N, Li Z, et al. Electrical pulse stimulation induces GLUT4 translocation in a Rac-Akt-dependent manner in C2C12 myotubes. FEBS Lett. 2018;592: 644–654.

27. Hamill KJ, Hopkinson SB, DeBiase P, Jones JC. BPAG1e maintains keratinocyte polarity through beta4 integrin-mediated modulation of Rac1 and cofilin activities. Mol Biol Cell. 2009;20: 2954–2962.

28. Wu J, Zhao R, Lin J, Liu B. Integrin β4 reduces DNA damagelilinduced p53 activation in colorectal cancer. Oncol Rep. 2018;40: 2183–2192.

29. Bierie B, Pierce SE, Kroeger C, et al. Integrin-β4 identifies cancer stem cell-enriched populations of partially mesenchymal carcinoma cells. Proc Natl Acad Sci U S A. 2017;114: E2337–e2346.

30. Ruan S, Lin M, Zhu Y, et al. Integrin β4-Targeted Cancer Immunotherapies Inhibit Tumor Growth and Decrease Metastasis. Cancer Res. 2020;80: 771–783.

31. Kang S, Jang Y, Chae YC, Kim BG.

32. Kim BG, Gao MQ, Choi YP, et al. Invasive breast cancer induces laminin-332 upregulation and integrin β4 neoexpression in myofibroblasts to confer an anoikis-resistant phenotype during tissue remodeling. Breast Cancer Res. 2012;14: R88.

33. Cherfils J, Zeghouf M. Regulation of small GTPases by GEFs, GAPs, and GDIs. Physiol Rev. 2013;93: 269–309.

34. Schmidt VA. Watch the GAP: Emerging Roles for IQ Motif-Containing GTPase-Activating Proteins IQGAPs in Hepatocellular Carcinoma. Int J Hepatol. 2012;2012: 958673.

35. Smith JM, Hedman AC, Sacks DB. IQGAPs choreograph cellular signaling from the membrane to the nucleus. Trends Cell Biol. 2015;25: 171–184.

36. Gorisse L, Li Z, Wagner CD, et al. Ubiquitination of the scaffold protein IQGAP1 diminishes its interaction with and activation of the Rho GTPase CDC42 2020;295: 4822–4835.

37. Woods D, Turchi JJ. Chemotherapy induced DNA damage response: convergence of drugs and pathways. Cancer Biol Ther. 2013;14: 379–389.

38. Cheung-Ong K, Giaever G, Nislow C. DNA-damaging agents in cancer chemotherapy: serendipity and chemical biology. Chem Biol. 2013;20: 648–659.

39. Kim C, Gao R, Sei E, et al. Chemoresistance Evolution in Triple-Negative Breast Cancer Delineated by Single-Cell Sequencing. Cell. 2018;173: 879–893.e813.

40. Noordermeer SM, van Attikum H. PARP Inhibitor Resistance: A Tug-of-War in BRCA-Mutated Cells. Trends Cell Biol. 2019;29: 820–834.

41. Geenen JJJ, Linn SC, Beijnen JH, Schellens JHM. PARP Inhibitors in the Treatment of Triple-Negative Breast Cancer. Clin Pharmacokinet. 2018;57: 427–437.

42. Lee A, Djamgoz MBA. Triple negative breast cancer: Emerging therapeutic modalities and novel combination therapies. Cancer Treat Rev. 2018;62: 110–122.

43. Campbell PS, Mavingire N, Khan S, et al. AhR ligand aminoflavone suppresses α6-integrin-Src-Akt signaling to attenuate tamoxifen resistance in breast cancer cells 2018;234: 108–121.

44. Ali HR, Dawson SJ, Blows FM, Provenzano E, Pharoah PD, Caldas C. Cancer stem cell markers in breast cancer: pathological, clinical and prognostic significance. Breast Cancer Res. 2011;13: R118.

45. Friedland JC, Lakins JN, Kazanietz MG, Chernoff J, Boettiger D, Weaver VM. alpha6beta4 integrin activates Rac-dependent p21-activated kinase 1 to drive NF-kappaB-dependent resistance to apoptosis in 3D mammary acini. J Cell Sci. 2007;120: 3700–3712.

46. Wei T, Lambert PF. Role of IQGAP1 in Carcinogenesis 2021;13.

47. Huang L, Xu S, Hu D, Lu W, Xie X, Cheng X. IQGAP1 Is Involved in Enhanced Aggressive Behavior of Epithelial Ovarian Cancer Stem Cell-Like Cells During Differentiation. Int J Gynecol Cancer. 2015;25: 559–565.

48. Deb S, Felix DA, Koch P. Tnfaip2/exoc3-driven lipid metabolism is essential for stem cell differentiation and organ homeostasis 2021;22: e49328.

49. Barutta F, Bellini S. Protective effect of the tunneling nanotube-TNFAIP2/M-sec system on podocyte autophagy in diabetic nephropathy 2022: 1–20.

50. Zhou L, Niu Z, Wang Y, et al. Senescence as a dictator of patient outcomes and therapeutic efficacies in human gastric cancer 2022;8: 13.

51. Guo F, Yuan Y. Tumor Necrosis Factor Alpha-Induced Proteins in Malignant Tumors: Progress and Prospects 2020;13: 3303–3318.

52. Liu Y, Ji X, Kang N, et al. Tumor necrosis factor α inhibition overcomes immunosuppressive M2b macrophage-induced bevacizumab resistance in triple-negative breast cancer. Cell Death Dis. 2020;11: 993.

53. Chen C, Sun X, Ran Q, et al. Ubiquitin-proteasome degradation of KLF5 transcription factor in cancer and untransformed epithelial cells. Oncogene. 2005;24: 3319–3327.

54. Chen C, Zhou Z, Ross JS, Zhou W, Dong JT. The amplified WWP1 gene is a potential molecular target in breast cancer. Int J Cancer. 2007;121: 80–87.

