## supplementary file 1 for "Integrin β4 promotes DNA damage-related drug resistance in triple-negative breast cancer via TNFAIP2/IQGAP1/RAC1"

中國科學院昆明動物研究所
KUNMING INSTITUTE OF ZOOLOGY
CHINESE ACADEMY OF SCIENCES

July 31, 2017

To whom it may concern,

Entrusted by Laboratory of The Kunming Institute of Zoology, Chinese Academy of Sciences, Kunming Cell Bank, Kunming Institute of Zoology, Chinese Academy of Sciences has conducted identification experiments on the Hcc1937 cell line ,and come to the following conclusions:

1. There was no third allele found in all the locations of Hcc1937 cell line ,it indicating that there was no cross-contaminant of human source cell line.
2. As the table shown below, compared the STR data of Hcc1937 cell line in the databases of ATCC,KCB and DSMZ, all the locations of Hcc1937 were exactly matched with the locations of Hcc1937 cells found in all the three cell banks, so it is Hcc1937 cell line.

| Gene Loci | Hcc1937 ( Sample ) | Hcc1937 ( KCB ) | Hcc1937 ( ATCC, DMSZ ) |
|---|---|---|---|
| Amelogenin | X | X | X |
| D7S820 | 9,10 | 9,10 | 9,10 |
| CSF1PO | 12 | 12 | 12 |
| TH01 | 6 | 6 | 6 |
| D13S317 | 13 | 13 | 13 |
| D16S539 | 13,14 | 13,14 | 13,14 |
| vWA | 16,17 | 16,17 | 16,17 |
| TPOX | 11 | 11 | 11 |
| D5S818 | 12 | 12 | 12 |

Best wishes

Wenhui Nie

Director
Kunming Cell Bank
Kunming Institute of Zoology, Chinese Academy of Sciences.
Kunming,China. 650223
Tel;+86 871 65195375


中國科學院昆明動物研究所
KUNMING INSTITUTE OF ZOOLOGY
CHINESE ACADEMY OF SCIENCES

July 31, 2017

To whom it may concern,

Entrusted by Laboratory of The Kunming Institute of Zoology, Chinese Academy of Sciences, Kunming Cell Bank, Kunming Institute of Zoology, Chinese Academy of Sciences has conducted identification experiments on the Hcc1806 cell line ,and come to the following conclusions:

1.There was no third allele found in all the locations of Hcc1806 cell line ,it indicating that there was no cross-contaminant of human source cell line.

2.As the table shown below, compared the STR data of Hcc1806 cell line in the databases of ATCC, DMSZ and KCB, all the locations of Hcc1806 were exactly matched with the locations of Hcc1806 cells found in all the three cell banks, so it is Hcc1806 cell line.

| Gene Loci | Hcc1806 ( Sample ) | Hcc1806 ( KCB ) | Hcc1806 ( ATCC, DMSZ ) |
|---|---|---|---|
| Amelogenin | X | X | X |
| D7S820 | 10,12 | 10,12 | 10,12 |
| CSF1PO | 12 | 12 | 12 |
| TH01 | 8 | 7,8 | 8 |
| D13S317 | 11 | 11 | 11 |
| D16S539 | 10 | 10 | 10 |
| vWA | 16,18 | 16,18 | 16,18 |
| TPOX | 8,9 | 8,9 | 8,9 |
| D5S818 | 13 | 13 | 13 |

Best wishes

Wenhui Nie

Director
Kunming Cell Bank
Kunming Institute of Zoology, Chinese Academy of Sciences.
Kunming,China. 650223
Tel;+86 871 65195375
