## supplementary file 2 for "Integrin β4 promotes DNA damage-related drug resistance in triple-negative breast cancer via TNFAIP2/IQGAP1/RAC1"

These cell lines (HCC1806, HCC1937, HEK293T) tested negative for mycoplasma contamination

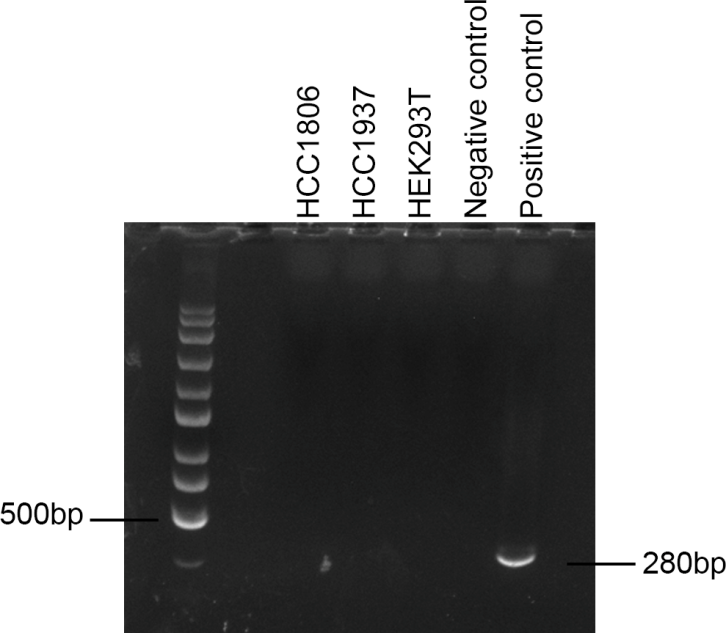

Protocol for detection of mycoplasma:

1. Collect culture medium for 2-3 days of cultured cells.

2. PCR system (Total 10 μl): Template (culture medium) 1ul, Upstream primer 0.5 μl (Forward primer: 5'-GGGAGCAAACAGGATTAGATACCCT-3'), Downstream primer 0.5 μl (Reverse primer: 5'-TGCACCATCTGTCACTCTGTTAACCTC-3'), 2 x Premix Ex Taq (CWBIO, CW0682M) 5 μl, ddH_2_O 3 μl.

3. PCR program : 94℃, 5 min, 94℃, 30 s, 30 cycles, 55℃, 30 s, 30 cycles, 72℃, 30 s, 30 cycles, 72℃ 5 min.

4. A PCR product of 280 bp indicates mycoplasma contamination.
