## supplementary file 3 for "Integrin β4 promotes DNA damage-related drug resistance in triple-negative breast cancer via TNFAIP2/IQGAP1/RAC1"

### 动物实验伦理审查意见书

### The Suggestion Of Animal Research Ethics

批准编号(ID Number) GDY2102096

<table>
<tr><td>实验名称<br>Project Title</td><td colspan="4">Integrin β4 通过 TNFAIP2 激活 Rac1 促进三阴性乳腺癌耐药研究</td></tr>
<tr><td>申请人姓名<br>Experimenter:</td><td colspan="4">崔秋霞</td></tr>
<tr><td>申请者单位<br>Project Organization:</td><td colspan="4">广东医科大学附属医院</td></tr>
<tr><td rowspan="4">使用动物情况<br>Situation of using laboratory animal</td><td colspan="4">动物来源 Source of laboratory animal: 北京维通利华实验动物技术有限公司</td></tr>
<tr><td>品种品系<br>Species or strain</td><td>小鼠</td><td colspan="2">等级 Grade SPF 级</td></tr>
<tr><td colspan="2">年龄 Age 6W-8W</td><td colspan="2">重量 Weight 12g-15g</td></tr>
<tr><td colspan="2">数量 Quantity 100 只 pcs<br>♀ 100 只 pcs ♂ 0 只 pcs</td><td colspan="2">实验日期 Experimental date:<br>2022.1.31-2024.2.1</td></tr>
<tr><td colspan="5">根据动物实验伦理原则，经审核本实验之实验动物选择、实验目的、实验方法、观测指标、实验结束后处死动物的方法等符合公认的 3R 原则（减少、替代、优化）原则，同意实施。Based on principles of animal research ethics, the experiment is approved for meeting the generally accepted 3R (Reduction, replacement, Refinement) principles such as the selection of laboratory animal, experimental purpose, methods, observation indicator, execution methods of experimental animals after experiments etc.</td></tr>
<tr><td>屏障设施条件<br>Condition of the barrier housing facility</td><td colspan="4">本设施的环境条件符合中国国家标准对屏障设施动物实验设施的有关标准，动物饲养管理和动物实验操作符合中国有关法规的要求。This housing facility is a barrier housing facility, and it's in keeping with national standard. The care of laboratory animal and the animal experimental operation have conforming to 《Administration Rule of Laboratory Animal》, et al.</td></tr>
<tr><td colspan="5">实验动物伦理委员会意见 Comments of laboratory animal ethical committee(LAEC)<br>根据本人申请，经评审，本课题符合相关伦理要求，同意以此申报材料申报相关课题。<br><br>签章 Stamp 广东医科大学实验动物伦理委员会 LAEC<br>日期 Date(yy/mm/dd): 2021-2-23 李华文</td></tr>
</table>

广东医科大学 实验动物伦理委员会 4408030012616
