## supplementary file 4 for "Integrin β4 promotes DNA damage-related drug resistance in triple-negative breast cancer via TNFAIP2/IQGAP1/RAC1"

### 伦理审查意见

| | | | |
|---|---|---|---|
| 受理号 | YS2021036 | 意见号 | YJYS2021036 |
| 项目名称 | Integrin β4 通过 TNFAIP2 激活 Rac1 促进三阴性乳腺癌耐药研究 | | |
| 项目来源 | 国家自然科学基金 | | |
| 研究单位 | 广东医科大学附属医院 | | |
| 主要研究者 | 崔秋霞 | | |
| 审查类别 | 预审查 | 审查方式 | 简易审查 |
| 审查日期 | 2021 年 3 月 1 日 | 审查地点 | NA |
| 审查委员 | 梁政 | | |
| 审查文件 | 课题标书 | | |
| 批准文件 | 课题标书 | | |

审查意见

根据卫生部《涉及人的生物医学研究伦理审查办法》(2016)、CFDA《药物临床试验质量管理规范(2003)》、《药物临床试验伦理审查工作指导原则》(2010年)、《医疗器械临床试验质量管理规范(2016)》、《人类遗传资源管理暂行办法(1998)》、《人类遗传资源采集、收集、买卖、出口、出境审批行政许可事项服务指南(2015)》、WMA《赫尔辛基宣言》和 CIOMS 《人体生物医学研究国际道德指南》的伦理原则，经本伦理委员会审查，意见如下：

1、该研究项目实验设计合理，有一定的先进性、科学性和实用性，有较好研究团队，可以完成计划，方案设计无重大伦理问题。同意申报国家自然基金项目。

2、请遵循赫尔辛基宣言、GCP 原则、遵循伦理委员会批准的方案开展临床研究，保护受试者的健康与权利。

3、项目立项后，请再次提出正式伦理审查，正式伦理审查批准通过方可启动项目。

| | |
|---|---|
| 主任委员签字 | 张[illegible] |
| 伦理委员会 | 广东医科大学附属医院机构审查伦理委员会 |
| 伦理委员会 GCP 声明<br>我院伦理委员组成及操作方式严格遵循 GCP（包括 ICH-GCP）及相关法律、法规的规定，实施各项操作规程。 | |
| 日期 | 2021 年 3 月 1 日 |
